# Altered histone modifications in *Aedes aegypti* following Rift Valley fever virus exposure

**DOI:** 10.1101/2025.09.11.675494

**Authors:** Hunter A. Ogg, Zoey M. Mikol, David C. King, Chad E. Mire, Zeyad Arhouma, Erin Osborne Nishimura, Rebekah C. Kading, Corey L Campbell

**Affiliations:** Center for Vectorborne Infectious Diseases, Colorado State University, Fort Collins, Colorado, 80523, USA; Department of Microbiology, Immunology, and Pathology, Colorado State University, Fort Collins, Colorado, 80523, USA; Department of Biochemistry and Molecular Biology, Colorado State University, Fort Collins, Colorado, 80523, USA; United States Department of Agriculture, Agricultural Research Services, National Bio and Agro-defense Facility, Foreign Arthropod-Borne Animal Diseases Research Unit, Manhattan, Kansas, USA

**Keywords:** Mosquito, vector competence, *Culicidae*, chromatin immunoprecipitation sequencing, differential expression, RNA-Seq, ChIP-Seq, histone modifications

## Abstract

When arthropod-borne viruses (arboviruses) are delivered to vector mosquitoes in an infectious bloodmeal, viral components interact with host proteins to hijack cells and initiate replication. The extent to which arbovirus infection alters mosquito host transcriptional and genomic regulatory processes is currently unknown. We hypothesized that histone modifications would be altered in mosquitoes exposed to Rift Valley fever virus (RVFV MP12, *Phlebovirus riftense*, family *Phleboviridae*). We interrogated transcriptome and chromatin landscapes in *Aedes aegypti* midguts by performing Cleavage Under Targets and Release Using Nuclease (CUT&RUN), using H3K27ac and H3K9me3 marks. Altered H3K27ac marks were identified following RVFV MP12 exposure, as well as upon bloodfeeding alone. It took several days for differential H3K27ac marks to be associated with differentially expressed genes (DEGs) in RVFV-exposed midguts. H3K27ac peaks showed progressive depletion as infection progressed. Gene set enrichment analysis revealed that immune response transcripts were enriched at 1 and 3 dpf (days post-feeding) but depleted by 7 dpf. Hedgehog/Gli (glioma-associated oncogene homolog) signaling pathway transcripts were depleted, indicating possible viral manipulation of cellular polarization. Moreover, at 7 dpf, 7 of 102 DEGs were proximal to differentially acetylated sites in a pattern expected to favor viral propagation. However, one transcript coding for an antiviral effector (LysM-TLDc domain protein) showed significant depletion of both H3K9me3 and H3K27ac marks. Analysis of midguts after a non-infectious bloodmeal versus sugar-fed controls revealed global changes to H3K27ac and H3K9me3 marks during and following the period of bloodmeal digestion. Differential H3K27ac marks were proximal to one quarter of all DEGs at 1 dpf, consistent with an important role of H3K27ac in bloodmeal digestion. These results demonstrate that H3K27ac and H3K9me3 patterns are altered upon virus exposure in a complex interplay that favors viral replication but is also countered by host responses to limit replication.

**Author Summary:** Mosquito-borne viruses produce a high disease burden across the globe. Though transgenic mosquitoes are being developed to help mitigate disease transmission, little is known of the ways in which arbovirus infection alters chromatin structure in mosquitoes. Arboviruses typically replicate and assemble on cytoplasmic membranes and have not been traditionally expected to alter processes in the nucleus. Some histone post-translational modifications, such as, e.g. histone 3 lysine 27 acetylation (H3K27ac), are associated with accessible chromatin regions, whereas others, e.g, histone 3 lysine 9 triple-methylation (H3K9me3), are associated with inaccessible chromatin. This work explores these chromatin modifications and the ways in which they are tied to gene expression changes. Multiple lines of evidence presented here support the hypothesis that H3K27ac levels change in RVFV-exposed mosquitoes and are associated with altered gene expression in a manner consistent with viral hijacking of gene expression in the nucleus. In addition, bloodfeeding alone also altered H3K27ac levels at 1-day post-feeding, and over one quarter of differentially expressed genes had changes to H3K27ac marks near transcription start sites, which is consistent with the idea that broad metabolic changes occur to support digestion and egg development.

## Introduction

Vector-borne disease outbreaks, particularly those caused by arthropod-borne viruses (arboviruses), have increased in frequency and intensity in recent years[1]. Mosquito-borne viruses are unique in that they must successfully replicate in an alternating fashion among invertebrate and vertebrate hosts. As intracellular pathogens, they interact with host proteins to hijack cellular processes and push metabolic activity in favor of replication and viral assembly.

Regardless of whether the host is a mammal or mosquito, selective repression of transcriptional activity occurs[2–4], which is a hallmark of cellular hijacking to enable efficient viral propagation. In vector mosquitoes, immune tolerance and resistance allow viral replication without pathological signs. Virus replication is affected by and, in turn, influences host genomic regulation (reviewed in [5]). Though histone modifications have been best studied with DNA viruses, recent evidence has also implicated histone modifications in the efficacy of RNA virus infection, including flavivirus infection of mosquitoes[6–9]. These alterations could underpin features of host tolerance and resistance in vectors.

Histone modifications occur during development [10] across taxa and are also important during the innate immune response[11, 12]. Histone 3 lysine 27 acetylation (H3K27ac) is a trademark of accessible chromatin, which facilitates entry and binding of proteins to initiate transcriptional activation. H3K27ac may also be a marker for enhancer activity and can be perturbed upon pathogen infection[13, 14]. In particular, H3K27ac levels are elevated during Zika virus infection in *Ae. aegypti* cell culture via acetyl transferase CBP activity[6]. In contrast, histone 3 lysine 9 triple-methylation (H3K9me3) is associated with heterochromatin or silenced genes (reviewed in [15]); though, during virus infection, specific genes may be derepressed [16]. Though the details of the overall structural features of mosquito chromatin are largely unknown[17], recent efforts have begun to explore distinct elements, such as the importance of histone modifications[18, 19]. Other studies used computational approaches to identify *cis* regulatory elements (CREs) that differed between dengue virus susceptible and resistant *Ae. aegypti* strains[20]. Still, much remains to be understood of the role of CREs and chromatin modifications in the vector response to arbovirus infection.

Here, we focus on Rift Valley fever virus (RVFV MP12, *Phlebovirus riftense*, family *Phleboviridae*), a vaccine strain for a zoonotic pathogen of concern in sub-saharan Africa[21, 22]. Our previous studies established *Ae. aegypti* as a suitable model for study of RVFV-mosquito interactions, including susceptibility to the MP12 vaccine strain[4, 23–25]. Previous vector competence studies led to the hypothesis that *Ae. aegypti* mounts an effective innate immune response against RVFV compared to the much more competent vector, *Culex tarsalis*, as evidenced by viral infection kinetics and differential expression of a key signaling gene, *dishevelled* [4, 23]. In nature, *Ae. aegypti* has expanded its range, which is of concern for spread of mosquito-borne viruses in the US and Europe[26–28].

To define the relationship between gene expression changes and genomic regulatory regions in arbovirus-infected mosquitoes, we chose to characterize H3K27ac and H3K9me3 histone modifications in midguts of RVFV-MP12-exposed *Ae. aegypti* adult females and compare them to gene expression(Fig 1). Midguts are often the first site of arbovirus replication in vector mosquitoes. Analysis of chromatin marks in mosquito tissues is made possible by recent advances in chromatin immunoprecipitation sequencing, specifically Cleavage Under Targets and Release Using Nuclease (CUT&RUN). This technique and similar approaches require substantially less biological material and have improved signal-to-noise, compared to traditional chromatin immunoprecipitation methods[29–31](Fig 1B). Midgut gene expression changes that occur following exposure to RVFV MP12 were paired with companion analyses of mosquitoes that had received a bloodmeal alone compared to sugar-fed controls. We followed up these studies with interrogation of H3K27ac and H3K9me3 signatures. Our initial hypothesis was bloodfeeding induces gene expression and associated chromatin modifications that set the metabolic foundation for successful arbovirus infection [32]. Moreover, we also predicted histone modifications would be altered following exposure to RVFV MP12 in a manner that favors viral hijacking of host processes, as evidenced by functional roles of differentially expressed genes. We focus first on the changes that occur following RVFV exposure and follow up with the control experiments.

**Figure 1.**
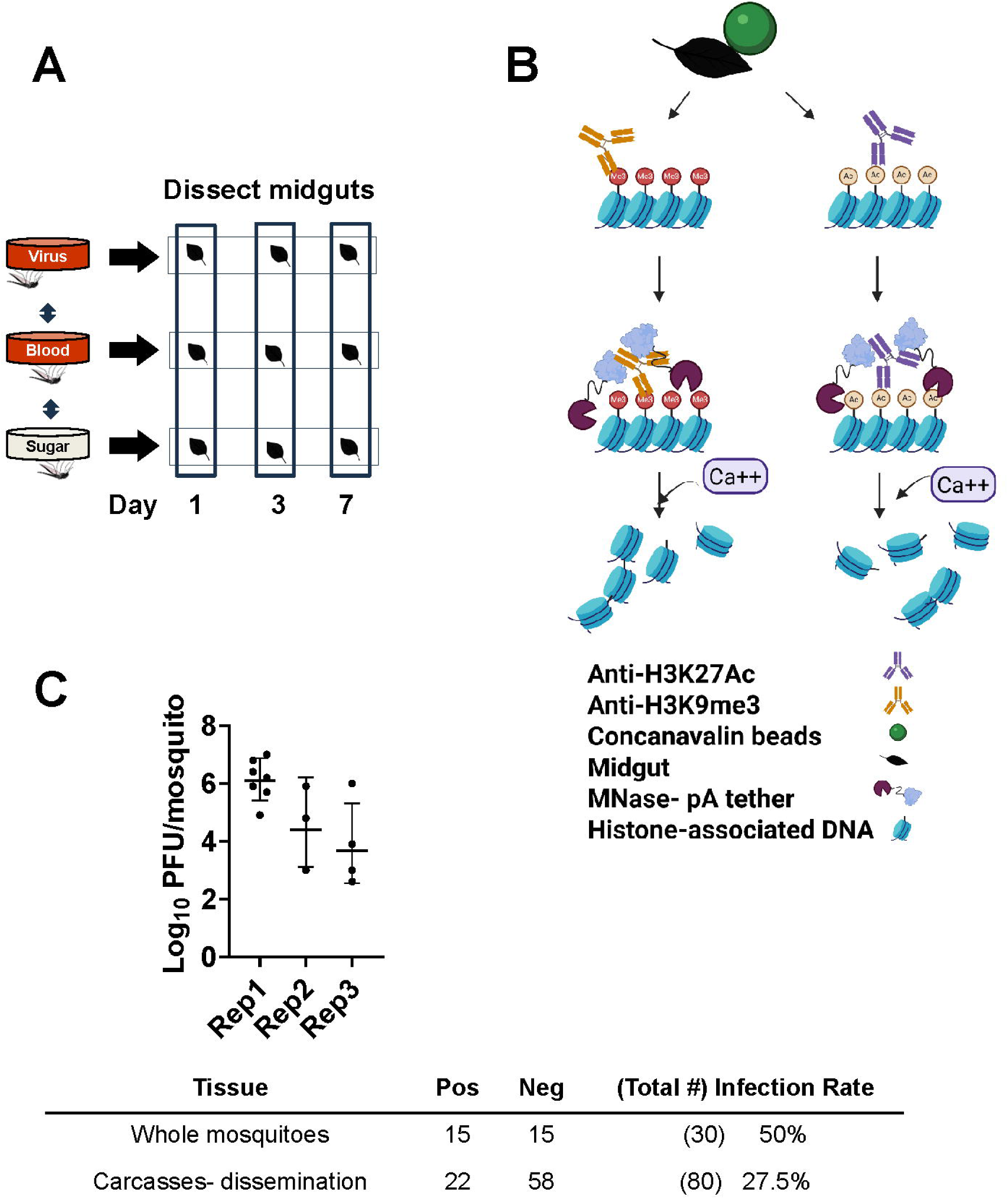
CUT&RUN enables capture of histone modifications. A. Two major experiments were done, RVFV MP12 vs bloodfed (RVFV v BF) or bloodfed vs sugar-fed (BF v SF) at 1, 3, or 7 dpf. Pools of 20 midguts were used for each sample in triplicate biological replicates of CUT&RUN or RNA-Seq. CUT&RUN libraries with inserts <55 nts were removed from analysis. DNA Peaks from merged libraries were called against input controls. B. General overview of CUT&RUN procedure shows that fixed midguts were bound to magnetic beads coated with concanavalin A (green), permeabilized with digitonin, and treated with antibodies specific for H3K27Ac or H3K9me3. Protein A/G-micrococcal nuclease fusion protein cleaves DNA at antibody binding sites for release into the supernatant, then subjected to purification and library preparation. Created in BioRender. (2026) https://BioRender.com/9y5ab72 C. Representative mosquito titers from RVFV-exposed whole mosquitoes collected at 7 dpf. Top row-Infection rates of mosquitoes shown in graph. Bottom row-Infection rates determined from representative carcasses following removal of midguts for CUT&RUN.

## Results

### Gene expression upon RVFV exposure

To gain insight into transcriptional profiles following RVFV-MP12 exposure, *Ae. aegypti* were provided an infectious oral bloodmeal, and pooled midguts were processed for RNA-Seq at 1, 3, and 7 days post-feeding (Fig 1A, S1 Table, S1 Fig). Viral bloodmeal titers were 6.9–7.8 log10 plaque-forming units (PFU) per ml. Virological confirmation showed that 50% of whole mosquitoes carried infectious virus at 7 days post-feeding (dpf), and a representative set of carcasses showed that about 28% of mosquitoes had disseminated infections (Fig 1C). For the RVFV v bloodfed (BF) comparison at 1 dpf, virus-exposed midgut pools had 94 differential expressed genes (DEGs, FDR <0.10, Fig 2, S2 Table) compared to blood-fed controls. Gene set enrichment analysis (GSEA) using our custom annotation was done to calculate the normalized enrichment score (NES) for each functional category across all the collection times (S3 Table)[33]. NES analysis indicated that overall signaling and immune response genes were enriched at 1 dpf, as was the broad category of replication, (DNA) repair, transcription, translation [34]. As the infection progressed, by 3 dpf, there were 2023 total DEGs. This functional category continued to be enriched, along with cytoskeletal and structural, immune and proteolytic pathways (Fig2B, S2 Fig, S3 Table). By 7 dpf, 102 DEGs were differentially expressed in the RVFV-exposed group, with 71 DEGs depleted, and just 31 transcripts enriched over BF controls (Fig 2, S2 Table). Custom GSEA analysis indicated that replication, (DNA) repair, transcription, translation transcript categories were still over-represented in virus-exposed midguts (S3 Table), which would be consistent with continued viral replication at this timepoint. However, immune response, proteolysis and metabolic functional categories were significantly under-represented, consistent with select transcriptional repression expected during viral hijacking[35, 36].

**Figure 2.**
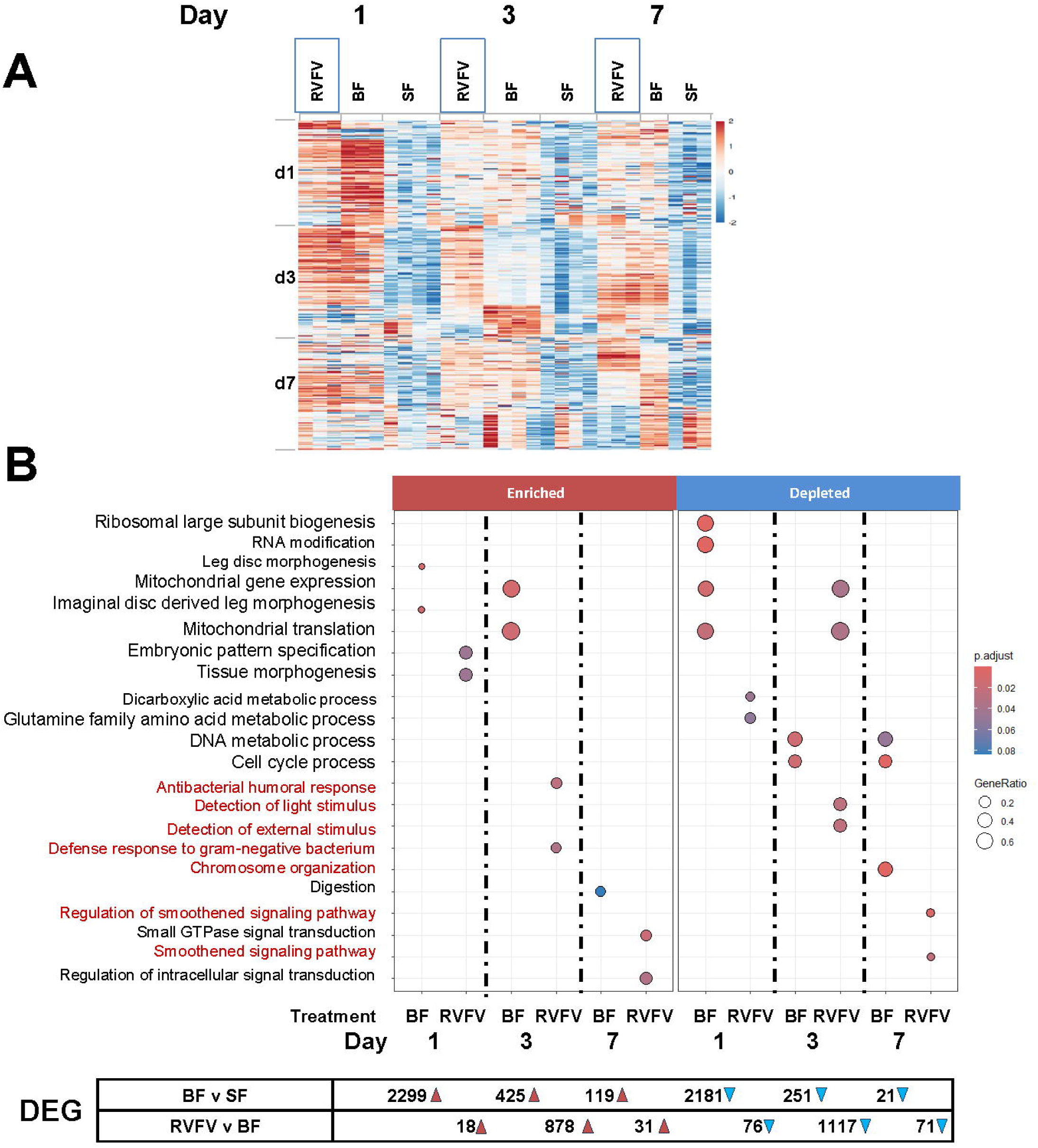
Differential gene expression profiles at 1, 3 and 7 days post-treatment. A. Heatmap of bulk RNA-Seq data pools of 20 midguts per timepoint were ranked by the top 100 genes of the RVFV treatment group on the indicated dpf. Columns group individual biological replicate z-scores of RPKM values for RVFV vs. BF by treatment group (RVFV, BF, SF), then by timepoint. Number of replicates ranged from 2 to 4. B. GO-GSEA dotplot of RNA-Seq differential expression categories for 1, 3 and 7 days post-treatment BF (bloodfed vs sugar-fed) and RVFV (RVFV-exposed vs BF). GeneRatio refers to the proportion of the gene set of interest divided by the total number of genes. Highlighted biological processes are noted in red font. Numbers below indicate DEGs that were enriched (red triangle) or depleted (blue triangle).

GSEA analyses are often limited by the number of genes in a transcriptome that are assigned functional terms. Because our custom annotation is limited in terms of scope and functional category, we applied gene ontology (GO) term GSEA (GO-GSEA) analysis based on eggNOG mapper orthological assignments [37]. GO-GSEA provided corroborative evidence that anti-microbial and signaling responses were enriched at 1 and 3 days following RVFV exposure (Fig 2B, S2 Fig). By 7 dpf, GTPase signaling and overall regulation of signaling processes were significantly enriched in RVFV exposed midguts compared to BF controls at 7dpf (Fig 2B, S2 Table). Lastly, constituents of the smoothened signaling pathway were depleted at 7 dpf but unaffected upon bloodfeeding alone. Smoothened is a signaling pathway associated with the Hedgehog/Gli signaling pathway, which is responsible for maintenance of cell polarity and tissue development[38]. To further support the role of the smoothened signaling pathway, the *Drosophila melanogaster* Ci/Gli consensus (GACCACCCA) was searched against the promoters of the *Ae. aegypti* genome. Overrepresentation analysis of 3221 unique genes across the genome with Ci/Gli promoters indicated that negative regulation of chromatin and epigenetic regulation as well as negative regulation of DNA-templated transcription were major processes controlled by this promoter(S3 Fig).

### Gene expression upon a noninfectious bloodmeal

To differentiate the effects of virus exposure from those of bloodfeeding alone, additional experiments were performed to reveal effects on differential gene expression and histone modifications. By providing proteinaceous nutrients critical for egg-laying, bloodfeeding is crucial for reproduction. Because arbovirus infection of mosquitoes intrinsically requires a bloodfeeding event, and it is therefore intricately tied to vector competence, we hypothesized that bloodfeeding alone modulates midgut gene expression to favor mosquito permissiveness to virus infection [32]. Comparison of blood-fed vs sugar-fed (BF v SF) mosquito midguts at 1 day post-feeding (dpf) revealed 4480 DEGs (FDR, <0.10, Fig 2, S1 and S4 Figs, S2 Table). The expression of digestive enzymes is well documented[39], cellular processes with the lowest adjusted p values (padj) at each timepoint are highlighted here. Multiple RNA processing and cellular biogenesis pathways were modulated 1 dpf. By 3 dpf, the bloodmeal was completely digested; nevertheless mitochondrial processes remained elevated (Fig 2B). By 7 dpf, processes affecting chromosome organization and cell cycle progression were modulated compared to sugar-fed controls (S4 Fig).

### Altered H3K27ac profiles in RVFV v BF

Next, to determine whether gene expression changes associated with bloodfeeding and RVFV exposure were tied to H3K27ac and H3K9me3 modifications, we performed CUT&RUN on midguts following a bloodmeal and at 1, 3, 7 dpf to RVFV MP12 (S4 and S5 Tables). RVFV exposure led to incremental changes to global H3K27ac patterns over time that were lower than that of BF at 1 dpf and gradually increased over the course of infection (Fig 3A, S5 Fig). By 7 dpf, RVFV-exposed midguts showed slightly higher levels of global acetylation patterns than BF controls (Fig 3A). Upon analysis of statistically significant differences, a striking pattern emerged. There were 31 significantly different H3K27ac peaks proximal to transcription start sites (TSS) at 1 dpf (Fig 3B). Of these, 24/30 (80%) peaks were depleted in RVFV-exposed and just 6 peaks showed higher levels than BF controls. Though the number of altered H3K27ac peaks were much greater following a bloodmeal alone, the overall proportion of depleted/enriched peaks were similar (74% peaks depleted).

**Figure 3.**
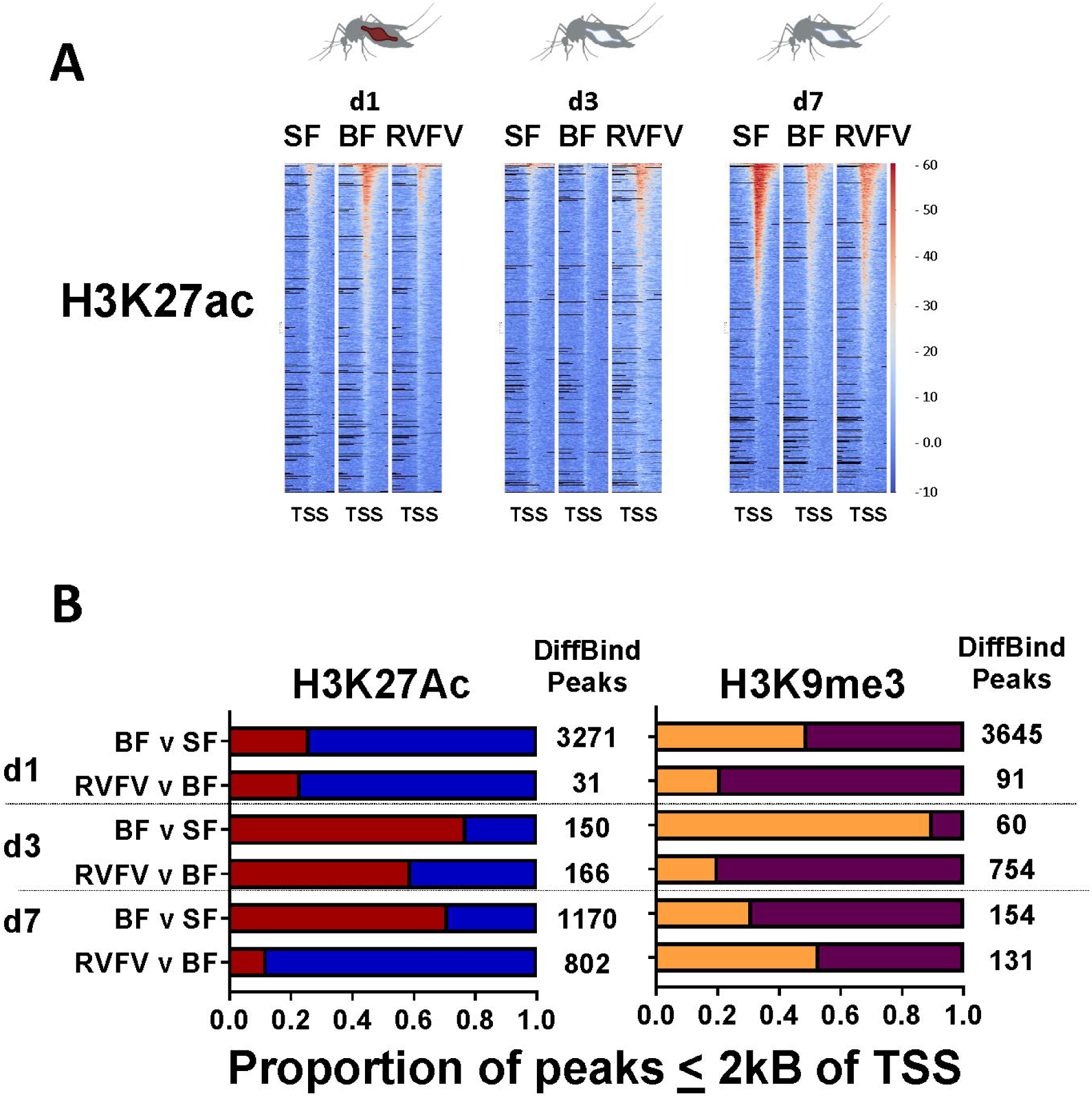
Midgut histone modification profiles. A. Representative H3K27ac peak heatmaps of aligned reads at 1, 3 and 7 days post-treatment show global trends within 2 kB of TSS (x axis). Reads were standardized to reads per genome coverage (RPGC). Y-axis indicates RPGC of input-subtracted read alignments. B. Proportion of DiffBind peaks within 2 kB of TSS (FDR <= 0.10) for H3K27ac, (red = enriched peaks compared to control, blue = depleted peaks compared to control) and H3K9me3 (orange = enriched, maroon = depleted). Total numbers of DiffBind peaks within 2 kB of TSS listed to the right of the bars.

By 3 dpf, an overall increase in H3K27ac enrichment occurred for both treatment groups (red bars, Fig 3B). Nevertheless, the proportion of significantly enriched H3K27ac peaks was reduced in RVFV-exposed (41%) compared to BF v SF treatment group(23% depleted, Fig 3B, S5 Table). This trend was even more dramatic at 7 dpf, wherein acetylation patterns in RVFV-exposed mosquitoes showed a marked reduction in acetylation sites compared to BF controls. In particular, 705/802 (88%) significant peaks were depleted in RVFV exposed midguts compared to just 29% in the BF v SF group (Fisher’s exact test, p<0.0001, Fig 3B, S5 Table). Genes of interest proximal to TSS in RVFV-treated samples showed trends in H3K27ac occupancy over time that are expected to affect cell morphogenesis at 1 dpf, regulation of transcriptional processes at 3 dpf and membrane transport by 7 dpf (S6 Fig). We currently cannot distinguish between the possibilities that overall dysregulation is occurring or whether these effects are due to viral hijacking.

Bloodfeeding alone was associated with changes to H3K27ac peak patterns that varied over the timecourse. Interestingly, global distributions of H3K27ac patterns increased upon bloodfeeding compared to SF but waned by 3 dpf (Fig 3A). Of 4073 quantitatively differential BFvSF CUT&RUN peaks at 1 dpf, 3271 were within 2 kB of DEG TSS (S4 Table, Fig3B). Of those, just 24% of peaks were quantitatively enriched in BF compared to SF controls. At 3 dpf, the overall numbers of differential peaks were lower, however the proportion of those that were enriched increased to 0.77. By 7 dpf, the sugar-fed group shows markedly higher levels of acetylation than either treatment group, which may be an indicator of aging in the absence of a bloodmeal (Fig 3A). GO-GSEA of genes proximal to TSS showed categories that were consistent with those of RNA-Seq data (S4 Fig), with genes affecting RNA metabolism showing modulation of H3K27ac marks at 1 dpf. These altered genomic regions supported changes to regulation of metabolic processes by 3 dpf. Finally, by 7 dpf, genes responsible for controlling chromosomal reorganization were identified proximal to H3K27ac sites.

Gene set over-representation analysis (GSORA) of gene functional categories for genes proximal to H3K27ac marks (MACS2 peaks) showed changes over time in blood-fed vs sugarfed controls(S7 Fig). In particular, at 1 dpf, peaks proximal to genes associated with chromosomal organization and histone modifications were altered in the BF treatment group, a pattern that shifted to catabolic and amide biosynthetic processes by 3 dpf (S7 Fig). These associations continued over time. In contrast, at 7 dpf, both SF and BF samples showed H3K27ac marks proximal to genes involved in several intracellular transport categories, including vesicle-mediated, amide and peptide transport (S7 Fig).

### RVFV DEGs with altered H3K27ac patterns

Correlation of DEGs with H3K27ac marks varied across the timecourse. In RVFV-exposed at 1 dpf, no differentially expressed genes had changes to proximal H3K27ac marks. However, at 3 dpf, 9 of 2023 DEGs did. By 7 dpf, there were 7/102 DEGs with depleted promoter-proximal acetylation patterns (S7 Table), 5 of which had enriched RNA levels, consistent with active gene expression. Of these 5 genes, 2 code for proteins that facilitate protein-protein interactions on cellular membranes, including AAEL005849 (synaptic vesicle protein) and AAEL019495 (pleckstrin/PDZ domain). Predicted gene function, coupled with transcript enrichment support the idea that these genes are proviral in supporting virus assembly or intracellular transport. A third gene, AAEL021746 (LysM-TLDc domain), which is predicted to regulate oxidative stress, showed enrichment of transcripts and depletion of H3K27ac marks[40]. The 2 genes with transcript depletion coupled with depletion of H3K27ac marks in RVFV-exposed midguts are predicted to have anti-viral function. One gene was a cytochrome P450 (CYP307A1, AAEL009762) and the second was a leucine-rich repeat (AAEL023746), which could be a novel pattern-recognition receptor.

### Post-bloodmeal gene expression changes coincide with histone modifications

A key goal of this study was to identify associations between gene expression changes and histone marks. One day following a noninfectious bloodmeal, about one fourth of all DEGs (1052/4480) showed proximal H3K27ac marks (S5 Table). There were 512 depleted DEGs among those that had altered peaks. Of those, 383/512 showed corroborative depletion of H3K27ac marks. However, for the enriched peaks, just 84/540 enriched DEGs showed the expected pattern of peak enrichment. S8 Fig shows the pattern of DEGs to promoter-proximal DiffBind peaks across all 3 chromosomes. This indicates that regulatory features cannot be explained by H3K27ac enrichment alone. At later timepoints following a bloodmeal alone, even fewer enriched DEGs showed corroborative H3K27ac enrichment, 6/676 and 11/140 at 3 and 7 dpf, respectively.

### Association between H3K9me3 peak patterns and RVFV-associated de-repression at 3 dpf

H3K9me3 marks may be associated with poised enhancers or transcriptional repression, depending on adjacent chromatin signatures[41, 42]. Our initial hypothesis was that H3K9me3 could underpin viral associated gene repression. Analysis of differentially bound H3K9me3 peaks by DiffBind showed depletion of peaks following RVFV exposure at 1 and 3 dpf (Fig 3B, maroon bar, S6 Table, S5 and S9 Figs), which was consistent with transcriptional de-repression. No DEGs overlapped with H3K9me3 peaks at 1 dpf. By 3 dpf, however, 109 DEGs showed peaks within 2 kB of TSS (S6 Table). Specifically, of 754 differentially bound peaks in the 3 dpf RVFV vs BF group, 109 DEGs showed proximal H3K9me3 peaks, 115 of 119 of these peaks were depleted. Peak depletion was corroborated by the observation that 60 of 109 DEGs were enriched (S2 Table). GO-GSEA analysis showed that processes affecting behavior and general metabolic processes were proximal to these marks. Therefore, instead of controlling part of viral repression of gene expression, as expected, our data was consistent with the idea that depletion of H3K9me3 peaks at 3 dpf was a correlate of gene de-repression.

By 7 dpf, just one DEG in the RVFV-exposed group, AAEL021746, showed significant depletion of H3K9me3 peaks. Overall, alterations in H3K9me3 marks were not associated with as many significant genes of interest as H3K27ac marks were, except for AAEL021746.

### Enhancers

Enhancers are long-range regulatory elements that activate or repress gene expression. H3K27ac marks are most often associated with enhancer activation of gene expression, whereas H3K9me3 enhancers are typically associated with transcriptionally repressed regions[42–44]. To explore potential involvement of enhancers, we interrogated our datasets for the presence of H3K27ac or H3K9me3 marks that were 50,000 to 200,000 nts away from TSSs. H3K27ac enhancers were associated with bloodfeeding at 1, 3 and 7 dpf (S10 Fig). At 1 dpf, the genes with these putative enhancer elements coded for RNA processing and metabolic changes (S10A Fig). In contrast, H3K27ac enhancers were enriched upon RVFV exposure only at 3 dpf.

Putative H3K9me3 enhancers were generally depleted upon bloodfeeding at 1 and 3 dpf (S10 FigB, maroon bar), which would be consistent with large-scale changes in metabolic needs, necessitating broad scale re-structuring of chromatin concomitant with gene expression changes. For RVFV exposed samples, enhancers were enriched at 1 and 7 dpf, which supports our hypothesis of virus-induced transcriptional repression. However, this interpretation is complicated by the lack of altered peaks at 3 dpf.

## Discussion

Global changes in H3K27ac marks occurred in a layered fashion in *Ae. aegypti* midguts upon receiving a non-infectious bloodmeal or concomitant RVFV MP12 exposure. At 1 dpf, BF v SF and RVFV v BF H3K27Ac peak profiles showed similarity in the overall numbers of peaks (Fig 3B). Blood-feeding alone led to an increase in enriched acetylation marks by 3 dpf, but those levels were reduced in RVFV-exposed midguts. By 7 dpf, RVFV exposed midguts showed significant depletion of H3K27ac marks relative to BF controls(Fig 3B). These results were corroborated in the RNA-Seq data, with 70% of DEGs being depleted in the RVFV-exposed group compared to BF controls at 7 dpf (Fig 2B). Moreover, the mean and median peak widths for peaks enriched in BF were significantly larger than those enriched for RVFV-exposed pools. Summary statistics of peak data indicate that RVFV-exposed pools had more peaks at promoter regions, whereas peaks in the BF treatment group had higher representation in gene bodies. These results are consistent with the idea that competitive regulatory processes are stimulated upon virus exposure, with the host activating gene expression to enhance anti-viral defense, while RVFV nonstructural protein on the S segment (NSs) or other viral protein(s) selectively alters gene expression to enhance infection [45, 46]. The patterns changed over time, indicating that elapsed time was required for significant histone modifications to occur. Alternatively, the increasing level of signal over time could be illustrative of a small percentage of infected cells at 1 dpf that expanded to a higher percentage over the course of infection[47].

The study of transcriptional regulation *in vivo* is critical, because histone modification patterns are not synonymous across cell culture and mosquito tissues [48]. Midguts are comprised of mixed cell populations with differing competency for virus infection[47]. The results captured here do not distinguish between different cell types or infection status. In addition, midgut pools were used for each replicate sample, thus reducing the high level of variability expected for individual mosquitoes[4]. Therefore, the patterns seen here represent overall trends associated with viral exposure.

RVFV-responsive host gene expression changes for a variety of functional categories occurred over time. For example, stimulation of cellular differentiation processes at 1 dpf were consistent with metabolic changes that might be required to support production of viral replication complexes(Fig 2C) [47]. These were coupled with repression of transcripts coding for dicarboxylic acid and glutamine metabolic processes, which could affect cellular signaling[49]. By 3 dpf, cellular processes specific to cell polarization processes were modulated (detection of light stimulus, phototransduction), as well as transcripts that control mitochondrial gene expression/translation (S3 Fig). By 7 dpf, signal transduction processes were repressed upon RVFV-exposure but not upon blood-feeding alone. In particular, components of the smoothened signaling pathway were depleted in the RVFV-exposed group(S4 Table, Fig3B). Smoothened is a frizzled class G-protein coupled receptor that acts in the hedgehog/Gli pathway to regulate cell polarity[38, 50]. Multiple lines of evidence indicate that arboviruses preferentially replicate in polarized cell types[51–53]. RVFV enters polarized mammalian cells more efficiently through the apical surface, and viral particles mature near basolateral membranes[54]. The smoothened pathway has antiviral characteristics in mammalian neuronal cells, therefore repression would likely favor viral propagation [55]. Though genes with C/Gli promoters were present across the sample sets (S8 Table), the gene set was not detected by GSEA until 7 dpf. We speculate that repression of transcripts in the smoothened pathway is proviral in mosquitoes; nevertheless, this will have to be explored further to determine the full extent of these interactions.

Interestingly, one DEG, AAEL021746 (LysM-TLDc domain), which was slightly enriched in RVFV exposed midguts at 7 dpf, showed significant depletion of both H3K9me3 and H3K27ac peaks. This gene also has a C/Gli promoter motif (S8 Table). LysM-TLDc containing proteins, e.g. human NCOA7 (Nuclear receptor coactivator 7), have been shown to interact with vacuolar-ATPase, which promotes endoplasmic vesicle acidification to impair infection of influenza A virus, human immunodeficiency virus and SARS-CoV[56–58]. This evidence suggests that LysM-TLDc domain containing proteins, such as AAEL021746, have anti-viral activity.

We demonstrated enrichment of immune response transcripts at 1 and 3 dpf, which were above and beyond what was induced by a noninfectious bloodmeal. RVFV-responsive genes of interest included defensin (AAEL003832, AAEL003857), cecropin (AAEL029046, AAEL029047), as well as transcriptional regulators that control immune transcriptional responses. However, by 7 dpf, immune response transcripts, e.g., pattern recognition proteins (AAEL001414, AAEL001420, AAEL010125, AAEL023746, AAEL024406) were depleted relative to BF controls.

Overall, these results are consistent with a gradual increase in the efficacy of viral repression of host gene expression over time. There are multiple examples of arboviral proteins that enter the nucleus to interact with transcriptional machinery[46, 59]. Bunyamwera (*Orthobunyavirus bunyamweraense)* nonstructural protein NSs interacts directly with *Mediator 8*[46]; the mediator complex ties RNA polymerase II to a host of transcription factors. In humans, the RVFV nonstructural NSs enters the nucleus to transcriptionally modulate beta catenin expression[45, 60]. NSs protein also suppresses host transcription, including of interferon beta, and degrades host protein kinase R (PKR, EIFAK2); these actions collectively facilitate viral replication as host cell machinery is co-opted and anti-viral responses are shut down[3, 61]. Even though the current study utilized an attenuated strain of RVFV, NSs remains fully functional[62]. In another study, *Ae. aegypti* infected with a deletion mutant strain of RVFV lacking NSs showed significantly reduced virus dissemination rates, indicating that this protein is playing some role in mosquito infection in addition to the roles described during vertebrate infection[24]. Therefore, progressive modulation of immune gene transcription and cell cycle processes in RVFV-infected mosquitoes may be related to NSs activity as an infection establishes in the mosquito midgut.

A previous report from Zhu et. al. showed that bloodfeeding alone prepared mosquitoes metabolically to favor arbovirus infection[32]. In the current study, over one quarter of all DEGs in the BF group at 1 dpf showed evidence of proximal H3K27ac marks. In addition, exposure to a noninfectious bloodmeal resulted in enrichment of immune response genes, likely a preventative measure to protect against potential pathogens present in the bloodmeal (S3 Table) [63]. Because H2K27ac promotes chromatin accessibility for gene expression, this result supports the hypothesis that bloodfeeding alone sets up favorable regulatory conditions that would favor viral hijacking. BF histone modification patterns were dramatically different than SF as early as 1 dpf, eased by 3 dpf and, then by 7 dpf, mosquitoes that had not received a bloodmeal showed substantially higher H3K27ac levels than the other treatment groups, consistent with a possible role in aging. Genes proximal to H3K27ac marks in SF mosquitoes did not change much over the course of time in terms of functional category type (S7 Fig).

A more competent vector might have less of a pronounced immune response at these early timepoints. *Ae. aegypti* has an intermediate level of competency for RVFV infection and transmission. Competence is much less than that of *Cx tarsalis* and higher than that of *Cx pipiens quinquefasciatus*[23, 64]. GO-GSEA analysis showed that defense processes were enriched at 1 and 3 dpf but repressed by 7 dpf. These observations were consistent with active immune response gene expression early in infection as part of overall intrinsic immune responses, which are key determinants of mosquito vector competence[65]. We would expect that a more susceptible mosquito species would show immune response repression earlier in the course of infection. However, further experiments are needed to sort out chromatin modulation across susceptible and resistant phenotypes.

Tri-methylation of histone 3 (H3K9me3) is associated with gene silencing. Due to the expected selective reduction in gene expression upon viral infection[45], we initially hypothesized that H3K9me3 marks might be enhanced during virus infection but found that instead, the global changes to H3K9me3 marks were much less dramatic than those found for H3K27ac. Nevertheless, significant depletion of H3K9me3 marks was observed in the 3 dpf RVFV vs BF group, which is consistent with derepression of gene expression. It is currently unclear whether this marked derepression at a time when viral replication complexes are active is part of viral hijacking or the host response.

Overall, these results provide support for the idea that histone modifications are part of the overall mosquito response to arbovirus infection. However, the extent to which specific changes are manipulated by viral proteins or a part of the host defense response must be determined by further experimentation.

## Conclusion

H3K27ac marks are significantly altered upon exposure to RVFV MP12, with a subset of DEGs that had significant changes indicative of active gene expression. However, competing processes for control of H3K27ac levels seem to be at play. Future efforts will continue to explore the epigenetic landscape layers in arbovirus exposed mosquitoes to identify the major players responsible for virus-induced transcriptional repression and whether these changes are consistent across arbovirus systems.

## Materials and Methods

### Mosquito and virus experiments

The Poza Rica strain of *Ae. aegypti* was colonized in 2012, originating from the state of Veracruz, Mexico[66]. Mosquito colonies were maintained at 28 °C on a 12:12 light:dark cycle; adults were fed water and sucrose (sugar cubes or raisins) *ad libitum*. Larvae were reared on TetraMin fish food (Spectrum Brands Pet, Blacksburg, VA, USA) ground in a coffee grinder.

Mosquitoes (3–7 days old) were starved for food and water for 24 hours prior to feeding, then orally exposed to RVFV MP-12 or conditioned cell culture media mixed 1:1 with defibrinated sheep blood, as described in Campbell et al.[23]. A high passage strain of MP12 (passage unknown, local lab passage 1), which was a gift from the US Department of Agriculture, was used for these experiments. In brief, freshly grown MP-12 RVFV was grown in Vero cells at a multiplicity of infection of 0.01 for 3 days and then mixed 1:1 in defibrinated calf blood (Colorado Serum Company, Denver, CO, USA), with 1 mM ATP as phagostimulant and orally provided to adult mosquitoes in water-jacketed feeders set to 37°C. Viral bloodmeal titers were at 6.9–7.8 log10 plaque-forming units (PFU) per ml. As a control, a separate group of age-matched mosquitoes were held for the indicated period. Three replicates of each experiment were performed.

Plaque assays were done using methods described previously[23]. Midgut infection rates were determined as follows. Individual mosquito carcasses were placed in 250 µl mosquito diluent (DMEM, 20% heat-inactivated FBS, 50 µg/ml Pen-Strep, 50 µg/ml gentamicin, and 2.5 µg/ml amphotericin B) following removal of midguts for CUT&RUN and stored at -80°C. Samples were homogenized on a Qiagen Tissuelyzer (Qiagen) at 30 beats per second frequency for 30 seconds. Homogenates were pelleted for 3 minutes at 21,000 x g, then 50 µl homogenate was placed into one well of a 12 well plate and assayed in duplicate. At 3 days post-infection, wells were assessed for cytopathic effects.

### CUT&RUN

Our approach followed that of Skene, et al.[30] with modifications. Specifically, pools of 20 midguts from blood-fed and RVFV MP12-fed *Aedes aegypti* were dissected 1, 3 and 7 days post feed or from sugar-fed controls. Midguts were placed into 200 µL of 1x Schneider’s Drosophila Media (<u>Gibco #21720024</u>). Following completion of dissections for all samples, 300 µL of formaldehyde fixation solution was added to each tube for a final concentration of 0.1% (using 16% methanol-free formaldehyde (Cell Signaling # 12606) in phosphate buffered saline (PBS) with 1x protease inhibitor cocktail (PIC, cOmplete™ Protease Inhibitor Cocktail, # 11697498001). After a 10-minute incubation at room temperature, 50 µL of 10x Glycine (<u>Cell Signaling #7005</u>) was added to stop the reactions and samples were incubated for another 5 minutes at room temperature (RT). Samples were centrifuged for 5 minutes at 2000xg 4°C then washed with 500 µL PIC/PBS. At all steps, tubes were flicked to mix solutions rather than pipetting to prevent loss of midguts. Midguts were resuspended in 1 ml Wash buffer (200mM HEPES pH 7.5, 1.5 M NaCl, 0.5 mM spermidine, 1x PIC), then centrifuged for 3 min at 2,000 xg at RT. Each reaction was resuspended in 100 µL 1x Wash buffer. Ten µL per sample of concanavalin A beads (Fisher # NC1831103) were activated using the methods of Skene, et al.[30]. Then the beads were added to the sample and incubated 5 minutes at RT. Each sample tube was placed on a magnetic stand (ThermoFisher 12321D). Once the solution cleared, the liquid was removed and replaced with 100 µL Digitonin solution (200mM HEPES pH 7.5, 1.5M NaCl, 20mM spermidine, 1x PIC, 0.1% digitonin) with 2mM EDTA and the appropriate antibody. The antibodies used in this study were 2 µg per reaction anti-H3K27ac Ab (Abcam, ab4729), or 1 µg per reaction anti-H3K9me3 Ab (Abcam, ab8898) and Rabbit (DA1E) mAb IgG XP® Isotype Control (Cell Signaling #66362) as a negative control. Samples were incubated overnight at 4°C on a rotator. Then the samples were put on the magnetic stand. Once the solution was clear, the samples were washed with 1 ml Digitonin buffer and put back on the stand. Upon removing the buffer, 50 µL solution containing 1.5 µL protein A/G-micrococcal nuclease (pAG/MNase, Epicypher, #15-1116) in Digitonin buffer was added per sample and mixed gently by flicking the tube. Samples were incubated at 4°C for 1 hour. The samples were then washed twice as before in 1 ml Digitonin buffer per sample. Samples were resuspended in 150 µL Digitonin buffer, placed on ice for 5 minutes, then activated by adding 3 µL 100 mM cold Calcium Chloride to each tube. Samples were incubated at 4°C for 30 minutes. Reactions were stopped by the addition of 150 µL Stop buffer (0.17 M NaCl, 10mM EDTA, 2mM EGTA, 0.1 mg/ml glycogen, 0.1% Digitonin, 8.3 mg/ml RNAse A). Samples were the centrifuged at 3 min at 16,000 xg for 2 minutes. Supernatants were transferred to new tubes containing 3 µL 10% sodium dodecyl sulfate and 2 µl Proteinase K (20 mg/ml). Samples were then incubated at 65°C for 2 hr, then transferred to low bind tubes. DNA fragments were purified using Qiagen PCR purification kits.

### Input sample preparation

Input samples were processed in parallel with the other samples through the first wash steps. Total DNA was extracted using a salt extraction method and resuspended in 100µl water [67]. Following extraction, samples were placed in a 0.65 mL Diagenode tube and sonicated in a Diagenode Bioruptor Pico (Diagenode) for 20 cycles of 30 seconds on, 30 seconds off. Fragment sizes of ∼200 bp were subsequently validated using an HS D1000 chip with an Agilent 4150 tapestation (Agilent).

### Sequencing

Three biological replicates were sequenced per sample group. Briefly, standard library preparation methods made use of the NEBNext® Ultra™ II DNA Library Prep Kit for Illumina (E7103L) following manufacturer’s directions with 1 ng DNA per library prep and ∼13 PCR cycles for the amplification step. Library quality was confirmed using high sensitivity DNA reagents (Q33231) on the Qubit 2 (Qubit) and Agilent tapestation 4150 (Agilent). Sequencing was performed on an Illumina NovaSeq (https://Azenta.com) with read depths indicated in S2 Table. Libraries with less than 55 nt inserts were removed from analysis (S2 Table).

### Analysis

All scripts utilized for the analysis of this data can be found on GitHub (https://github.com/CRosenbergCode/Aedes_aegypti_Midgut_CUT-RUN), Sequencing reads were trimmed and filtered using Fastp version 0.23.4[68]. Duplicate reads were removed for ChIP-seq reads but retained for RNA-seq analysis[69]. These trimmed and filtered reads were aligned to an index prepared from the VectorBase-68_AaegyptiLVP_AGWG_Genome.fasta using HISAT2 version 2.2.1[70, 71]. Only paired reads which aligned concordantly were retained. Data regarding alignment rates and quality parameters per sample can be found in Supplemental Tables 1-3. Peak calling was performed on merged alignments using MACS2 macs2 2.2.9.1 using --keep dups all, qvalue 0.05 against VectorBase-68_AaegyptiLVP_AGWG_Genome.fasta [71, 72]. All samples were standardized using the corresponding input sample.

Bigwig files were created using Deeptools bamCoverage with a bin size of 10 bases and normalized using reads per genome coverage (RPGC)[73]. They were then compared against their corresponding inputs using Deeptools bigwigCompare, with a bin size of 10 bases and -- operation subtract, thus generating bigwigs that use direct subtraction as opposed to fold change. Peak heatmaps and read density maps were made using deepTools functions “computeMatrix”, and “plotHeatmap”. computeMatrix was run with the “make_matrices.sh” script using VectorBase-68_AaegyptiLVP_AGWG_Genome.bed (No MIT, long-noncoding RNAs or pseudogenes) as reference.

Differential binding analysis was performed using Diffbind version 3.12.0 with the edgeR option [74] in R version 4.3.2[75] using a false discovery rate(FDR) threshold of p<0.10. A custom script was created to determine the proximity of DiffBind peaks to genes or other features of interest in the *Ae. aegypti* genome (S4 and S5 Tables).

### RNA Sequencing and analysis

Three to 4 biological replicates were sequenced per sample group. Bulk RNA-Seq libraries were prepared from 200 ng total RNA, using polyA+ purification (NEBNext® Poly(A) mRNA Magnetic Isolation Module) prior to library prep with the NEBNext® Ultra™ II Directional RNA Library Prep Kit for Illumina. Sequencing was performed on an Illumina NovaSeq (https://Azenta.com) with read depths indicated in S1 Table.

For RNA-seq alignment, a reference transcriptome was created using VectorBase-68_AaegyptiLVP_AGWG_AnnotatedTranscripts.fasta [71] using HISAT2 (2.2.1)[70], then alignment was performed using standard parameters. Gene-level aggregation was performed using HTseq-count (2.0.5). Differential expression analysis was performed using DEseq2 (1.48.0).

### Gene Set Enrichment analysis

GO-GSEA libraries were built using AnnotationForge version 2.13 in R. Genome-wide annotation was performed using Entrez gene identifiers[39]. The fast gene set enrichment analysis algorithm R implemented by the fgsea R package (version 1.28.0) [76] was used for GO-GSEA analysis[77]. GO terms were obtained using EggNog Mapper 2.1.12, limiting the orthology search to Diptera only[37]. Custom annotations were utilized to define broader functional categories as described in Campbell et al 2019 [33]. For both GSEA (ranked nonparametric analysis) and ORA (simple arithmetical calculation that does not involve ranking), a minimum size of 15 and a maximum size of 500 were required for a GO annotation to be included in analysis. Ranking was based on the -log10(p-value). In all cases, ordering was decided arbitrarily in case of ties. GO-GSEA of RNA-seq differential binding was based on DESeq2 results. GO-GSEA of ChIP-seq differential binding was based on Diffbind results. In cases of multiple peaks corresponding to the same gene, only the peak with the highest p-value was retained. For ORA, duplicate peaks for the same gene were removed to ensure each gene only occurred once. For analysis of promoters, only peaks within 2000 bp of the TSS were included for the analysis. For analysis of enhancers, only peaks between 50,000 and 200,000 bp away from a TSS that were annotated as distal using ChIPSeeker were included for the analysis[78, 79]. The custom annotation was manually compiled based on Campbell et al 2019, which relied on VectorBase orthological assignments[33, 71].The custom OrgDB database and GMT files used are provided on the github for this manuscript. Visualization of gene enrichment utilized enrichplot version 1.28 [80] and ClusterProfiler version 4.15[81].

### Motif Enrichment analysis

Enrichment analysis was performed using HOMER (Hypergeometric Optimization of Motif EnRichment) version 5.1 with a custom genome created using VectorBase-68_AaegyptiLVP_AGWG annotations[82]. An offset of 2000 base pairs upstream and 200 base pairs downstream of the transcription start site were used and a random selection of promoter regions was used as the background. For searching known motifs such as CI/GLI binding sites, the consensus sequence (GACCACCCA) in *D. melanogaster* was used and a maximum of 1 mismatch was allowed. Genes from this study with these motifs are in S8 Table.

## Supporting Information

**S1 Table.** RNA-Seq raw reads with Q20/Q30 scores, insert sizes and alignment rates.

**S2 Table.** RNA-Seq DESeq2 analysis for all samples and timepoints.

**S3 Table.** Normalized Enrichment Score Table for RVFV v BF time course.

**S4 Table.** CUT&RUN-Seq raw reads with Q20/Q30 scores, insert sizes and alignment rates.

**S5 Table.** H3K27ac DiffBind output for genes within 2kB of TSS.

**S6 Table.** H3K9me3 DiffBind output for genes within 2kB of TSS.

**S7 Table.** RVFV v BF DEGs with proximal H3K27Ac peaks at the 7dpe

**S8 Table.** CI/GLI promoter regions identified using HOMER.

**S1 Fig. MA plots of RNA-Seq differential expression data**.

**S2 Fig. GO-GSEA of RVFV v BF RNA-Seq differential expression data.**

**S3 Fig. Over-representation analysis of all *Ae. aegypti* genes with Ci/Gli motif.**

**S4 Fig. GO-GSEA of BF vs SF RNA-Seq transcript differential expression data.**

**S5 Fig. H3K27ac and H3K9me3 peak profiles.** Top row: H3K27ac Input-subtracted peak heatmaps of aligned reads at 1, 3 and 7 days post-treatment show global trends within 2 kB of TSS (x axis). Y-axis indicates RPGC (reads per genome coverage) of input-subtracted read alignments. Notice the different Y-axis scales. Bottom row: H3K9me3 Input-subtracted peak heatmaps.

**S6 Fig. RVFV vs BF datasets: GSEA of GOIs proximal to H3K27Ac and H3K9me3 marks.**

**S7 Fig. GO categories for genes proximal to H3K27ac marks change over time.** A. GO terms for genes within 2kB of TSS (MACS2 peak calls) displayed using over-representation analysis. X axis shows collection day (d1, d3, or d7), BF, bloodfed, SF, sugar-fed. Number in parentheses below the sample name indicates the number of macs2 peaks considered in the analysis. This is a qualitative analysis and does not indicate statistically significant differences between groups. B. Transition of GO terms over time in BF midguts. GO terms for genes within 2kB of TSS (DiffBind peak calls) displayed using over-representation analysis.

**S8 Fig. BFvSF 1 dpf: Relationship of H3K27ac peaks and DEGs.** Coordinates for DEG start sites (DEG Log2FC, right y-axis) were plotted alongside H3K27ac peak coordinated and fold-change values (Peak FC, left y-axis).

**S9 Fig. H3K9me3 RVFV vs BF vs SF peak heatmaps.** Input-subtracted H3K9me3 peak heatmaps of aligned reads at 1, 3 and 7 days post-treatment show global trends within 2 kB of TSS (x axis). Y-axis indicates RPGC (reads per genome coverage) of input-subtracted read alignments.

**S10 Fig. Enhancers.** A. GSEA functional groups of genes at the BF v SF 1 dpf timepoint that showed predicted H3K27ac peaks. B. Proportion of DiffBind peaks within 2 kB of TSS (FDR <= 0.10) for H3K27ac, (red = enriched, blue = depleted) and H3K9me3 (orange = enriched, maroon = depleted). Total numbers of DiffBind peaks within 2 kB of TSS listed to the right of the bars.

## Acknowledgements

We thank the Swygert lab at Colorado State University for the use of their Bioruptor Pico instrument. RVFV MP-12 virus stock was kindly provided by the United States Department of Agriculture, Agricultural Research Service. Figures 1A and 1B were generated in BioRender (https://BioRender.com).

## Informed Consent Statement

Not applicable.

## Data Availability Statement

Raw data available upon request. All raw sequencing data has been uploaded to the NCBI Sequence Read archive under Bioprojects PRJNA1284329 and PRJNA1290393.

## Author Contributions

**Conceptualization:** C Campbell

**Data curation:** C Campbell, H Ogg

**Formal analysis:** Z Arhouma, C Campbell, H Ogg, Z Mikol

**Funding acquisition:** C Campbell, C Mire

**Methodology:** Z Arhouma, D King, H Ogg, Z Mikol

**Validation:** E Osborne Nishimura, D King, H Ogg, C Campbell

**Project administration:** C Campbell

**Visualization:** Z Mikol, H Ogg

**Resources:** R Kading, C Mire, C Campbell

**Writing, original:** C Campbell, H Ogg

**Writing, review and editing:** C Campbell, C Mire, R Kading, E Osborne Nishimura, D King

## References

1. Weaver SC, Charlier C, Vasilakis N, and Lecuit M, Zika, Chikungunya, and Other Emerging Vector-Borne Viral Diseases. Annu Rev Med, 2018. 69: p. 395–408. DOI: 10.1146/annurev-med-050715-105122.

2. Billecocq A, Spiegel M, Vialat P, Kohl A, Weber F, Bouloy M, et al., NSs protein of Rift Valley fever virus blocks interferon production by inhibiting host gene transcription. J Virol, 2004. 78(18): p. 9798–806. DOI: 10.1128/JVI.78.18.9798-9806.2004.

3. Ikegami T, Narayanan K, Won S, Kamitani W, Peters CJ, and Makino S, Rift Valley fever virus NSs protein promotes post-transcriptional downregulation of protein kinase PKR and inhibits eIF2alpha phosphorylation. PLoS Pathog, 2009. 5(2): p. e1000287. DOI: 10.1371/journal.ppat.1000287.

4. Smith CB, Hodges NF, Kading RC, and Campbell CL, Dishevelled Has Anti-Viral Activity in Rift Valley Fever Virus Infected Aedes aegypti. Viruses, 2023. 15(11). DOI: 10.3390/v15112140.

5. Rehman UU, Ghafoor D, Ullah A, Ahmad R, and Hanif S, Epigenetics regulation during virus-host interaction and their effects on the virus and host cell. Microb Pathog, 2023. 182: p. 106271. DOI: 10.1016/j.micpath.2023.106271.

6. Amarante AM, da Silva ICA, Carneiro VC, Vicentino ARR, Pinto MA, Higa LM, et al., Zika virus infection drives epigenetic modulation of immunity by the histone acetyltransferase CBP of Aedes aegypti. PLoS Negl Trop Dis, 2022. 16(6): p. e0010559. DOI: 10.1371/journal.pntd.0010559.

7. Caraballo GI, Rosales R, Viettri M, Castillo JM, Cruz R, Ding S, et al., The Dengue Virus Nonstructural Protein 1 (NS1) Interacts with the Putative Epigenetic Regulator DIDO1 to Promote Flavivirus Replication in Mosquito Cells. J Virol, 2022. 96(12): p. e0070422. DOI: 10.1128/jvi.00704-22.

8. Harrell T, Basak S, Sultana H, and Neelakanta G, Zika virus modulates arthropod histone methylation for its survival in mosquito cells. PLoS One, 2025. 20(2): p. e0319290. DOI: 10.1371/journal.pone.0319290.

9. Zhou D, Wu Z, Park JG, Fiches GN, Li TW, Ma Q, et al., FACT subunit SUPT16H associates with BRD4 and contributes to silencing of interferon signaling. Nucleic Acids Res, 2022. 50(15): p. 8700–8718. DOI: 10.1093/nar/gkac645.

10. Gaddelapati SC, Albishi NM, Dhandapani RK, and Palli SR, Juvenile hormone-induced histone deacetylase 3 suppresses apoptosis to maintain larval midgut in the yellow fever mosquito. Proc Natl Acad Sci U S A, 2022. 119(11): p. e2118871119. DOI: 10.1073/pnas.2118871119.

11. Carson WFt and Kunkel SL, Regulation of Cellular Immune Responses in Sepsis by Histone Modifications. Adv Protein Chem Struct Biol, 2017. 106: p. 191–225. DOI: 10.1016/bs.apcsb.2016.08.004.

12. Daskalaki MG, Tsatsanis C, and Kampranis SC, Histone methylation and acetylation in macrophages as a mechanism for regulation of inflammatory responses. J Cell Physiol, 2018. 233(9): p. 6495–6507. DOI: 10.1002/jcp.26497.

13. Chen CH, Zheng R, Tokheim C, Dong X, Fan J, Wan C, et al., Determinants of transcription factor regulatory range. Nat Commun, 2020. 11(1): p. 2472. DOI: 10.1038/s41467-020-16106-x.

14. Del Rosario RCH, Poschmann J, Lim C, Cheng CY, Kumar P, Riou C, et al., Histone acetylome-wide associations in immune cells from individuals with active Mycobacterium tuberculosis infection. Nat Microbiol, 2022. 7(2): p. 312–326. DOI: 10.1038/s41564-021-01049-w.

15. Padeken J, Methot SP, and Gasser SM, Establishment of H3K9-methylated heterochromatin and its functions in tissue differentiation and maintenance. Nat Rev Mol Cell Biol, 2022. 23(9): p. 623–640. DOI: 10.1038/s41580-022-00483-w.

16. Li F, Nellaker C, Sabunciyan S, Yolken RH, Jones-Brando L, Johansson AS, et al., Transcriptional derepression of the ERVWE1 locus following influenza A virus infection. J Virol, 2014. 88(8): p. 4328–37. DOI: 10.1128/JVI.03628-13.

17. Lezcano OM, Sanchez-Polo M, Ruiz JL, and Gomez-Diaz E, Chromatin Structure and Function in Mosquitoes. Front Genet, 2020. 11: p. 602949. DOI: 10.3389/fgene.2020.602949.

18. Gomez-Diaz E, Rivero A, Chandre F, and Corces VG, Insights into the epigenomic landscape of the human malaria vector Anopheles gambiae. Front Genet, 2014. 5: p. 277. DOI: 10.3389/fgene.2014.00277.

19. Ruiz JL, Yerbanga RS, Lefevre T, Ouedraogo JB, Corces VG, and Gomez-Diaz E, Chromatin changes in Anopheles gambiae induced by Plasmodium falciparum infection. Epigenetics Chromatin, 2019. 12(1): p. 5. DOI: 10.1186/s13072-018-0250-9.

20. Behura SK, Sarro J, Li P, Mysore K, Severson DW, Emrich SJ, et al., High-throughput cis-regulatory element discovery in the vector mosquito Aedes aegypti. BMC Genomics, 2016. 17: p. 341. DOI: 10.1186/s12864-016-2468-x.

21. Clark MHA, Warimwe GM, Di Nardo A, Lyons NA, and Gubbins S, Systematic literature review of Rift Valley fever virus seroprevalence in livestock, wildlife and humans in Africa from 1968 to 2016. PLoS Negl Trop Dis, 2018. 12(7): p. e0006627. DOI: 10.1371/journal.pntd.0006627.

22. Sang R, Kioko E, Lutomiah J, Warigia M, Ochieng C, O’Guinn M, et al., Rift Valley fever virus epidemic in Kenya, 2006/2007: the entomologic investigations. Am J Trop Med Hyg, 2010. 83(2 Suppl): p. 28-37. DOI: 10.4269/ajtmh.2010.09-0319.

23. Campbell CL, Snell TK, Bennett S, Wyckoff JH, 3rd, Heaslip D, Flatt J, et al., Safety study of Rift Valley Fever human vaccine candidate (DDVax) in mosquitoes. Transbound Emerg Dis, 2022. 69(5): p. 2621–2633. DOI: 10.1111/tbed.14415.

24. Crabtree MB, Kent Crockett RJ, Bird BH, Nichol ST, Erickson BR, Biggerstaff BJ, et al., Infection and transmission of Rift Valley fever viruses lacking the NSs and/or NSm genes in mosquitoes: potential role for NSm in mosquito infection. PLoS Negl Trop Dis, 2012. 6(5): p. e1639. DOI: 10.1371/journal.pntd.0001639.

25. Kading RC, Crabtree MB, Bird BH, Nichol ST, Erickson BR, Horiuchi K, et al., Deletion of the NSm virulence gene of Rift Valley fever virus inhibits virus replication in and dissemination from the midgut of Aedes aegypti mosquitoes. PLoS Negl Trop Dis, 2014. 8(2): p. e2670. DOI: 10.1371/journal.pntd.0002670.

26. Laporta GZ, Potter AM, Oliveira JFA, Bourke BP, Pecor DB, and Linton YM, Global Distribution of Aedes aegypti and Aedes albopictus in a Climate Change Scenario of Regional Rivalry. Insects, 2023. 14(1). DOI: 10.3390/insects14010049.

27. O’Dell N, Bolling BG, Dacko N, Carr JT, Hambrick B, Chaves LF, et al., Identifying environmental drivers of Aedes aegypti and Aedes albopictus abundance in the Dallas-Fort Worth metroplex using Random Forest modeling. J Med Entomol, 2025. DOI: 10.1093/jme/tjaf036.

28. Parker C, Ramirez D, and Connelly CR, State-wide survey of Aedes aegypti and Aedes albopictus (Diptera: Culicidae) in Florida. J Vector Ecol, 2019. 44(2): p. 210–215. DOI: 10.1111/jvec.12351.

29. Kaya-Okur HS, Janssens DH, Henikoff JG, Ahmad K, and Henikoff S, Efficient low-cost chromatin profiling with CUT&Tag. Nat Protoc, 2020. 15(10): p. 3264–3283. DOI: 10.1038/s41596-020-0373-x.

30. Skene PJ, Henikoff JG, and Henikoff S, Targeted in situ genome-wide profiling with high efficiency for low cell numbers. Nat Protoc, 2018. 13(5): p. 1006–1019. DOI: 10.1038/nprot.2018.015.

31. Yin H, Sweeney S, Raha D, Snyder M, and Lin H, A high-resolution whole-genome map of key chromatin modifications in the adult *Drosophila melanogaster*. PLoS Genet, 2011. 7(12): p. e1002380. DOI: 10.1371/journal.pgen.1002380.

32. Zhu Y, Zhang R, Zhang B, Zhao T, Wang P, Liang G, et al., Blood meal acquisition enhances arbovirus replication in mosquitoes through activation of the GABAergic system. Nat Commun, 2017. 8(1): p. 1262. DOI: 10.1038/s41467-017-01244-6.

33. Campbell CL, Saavedra-Rodriguez K, Kubik TD, Lenhart A, Lozano-Fuentes S, and Black WCI, Vgsc-interacting proteins are genetically associated with pyrethroid resistance in *Aedes aegypti*. PLoS One, 2019. 14(1): p. e0211497. DOI: 10.1371/journal.pone.0211497.

34. Kubik TD, Snell TK, Saavedra-Rodriguez K, Wilusz J, Anderson JR, Lozano-Fuentes S, et al., Aedes aegypti miRNA-33 modulates permethrin induced toxicity by regulating VGSC transcripts. Sci Rep, 2021. 11(1): p. 7301. DOI: 10.1038/s41598-021-86665-6.

35. Ma Y, Liang Y, Wang N, Cui L, Chen Z, Wu H, et al., Avian Flavivirus Infection of Monocytes/Macrophages by Extensive Subversion of Host Antiviral Innate Immune Responses. J Virol, 2019. 93(22). DOI: 10.1128/JVI.00978-19.

36. Pott F, Postmus D, Brown RJP, Wyler E, Neumann E, Landthaler M, et al., Single-cell analysis of arthritogenic alphavirus-infected human synovial fibroblasts links low abundance of viral RNA to induction of innate immunity and arthralgia-associated gene expression. Emerg Microbes Infect, 2021. 10(1): p. 2151–2168. DOI: 10.1080/22221751.2021.2000891.

37. Cantalapiedra CP, Hernandez-Plaza A, Letunic I, Bork P, and Huerta-Cepas J, eggNOG-mapper v2: Functional Annotation, Orthology Assignments, and Domain Prediction at the Metagenomic Scale. Mol Biol Evol, 2021. 38(12): p. 5825–5829. DOI: 10.1093/molbev/msab293.

38. Nusslein-Volhard C and Wieschaus E, Mutations affecting segment number and polarity in Drosophila. Nature, 1980. 287(5785): p. 795–801. DOI: 10.1038/287795a0.

39. Hixson B, Bing XL, Yang X, Bonfini A, Nagy P, and Buchon N, A transcriptomic atlas of Aedes aegypti reveals detailed functional organization of major body parts and gut regional specializations in sugar-fed and blood-fed adult females. Elife, 2022. 11. DOI: 10.7554/eLife.76132.

40. Jaramillo-Gutierrez G, Molina-Cruz A, Kumar S, and Barillas-Mury C, The Anopheles gambiae oxidation resistance 1 (OXR1) gene regulates expression of enzymes that detoxify reactive oxygen species. PLoS One, 2010. 5(6): p. e11168. DOI: 10.1371/journal.pone.0011168.

41. Blackledge NP and Klose RJ, The molecular principles of gene regulation by Polycomb repressive complexes. Nat Rev Mol Cell Biol, 2021. 22(12): p. 815–833. DOI: 10.1038/s41580-021-00398-y.

42. Zentner GE, Tesar PJ, and Scacheri PC, Epigenetic signatures distinguish multiple classes of enhancers with distinct cellular functions. Genome Res, 2011. 21(8): p. 1273–83. DOI: 10.1101/gr.122382.111.

43. Creyghton MP, Cheng AW, Welstead GG, Kooistra T, Carey BW, Steine EJ, et al., Histone H3K27ac separates active from poised enhancers and predicts developmental state. Proc Natl Acad Sci U S A, 2010. 107(50): p. 21931–6. DOI: 10.1073/pnas.1016071107.

44. Zhu Y, van Essen D, and Saccani S, Cell-type-specific control of enhancer activity by H3K9 trimethylation. Mol Cell, 2012. 46(4): p. 408–23. DOI: 10.1016/j.molcel.2012.05.011.

45. Le May N, Mansuroglu Z, Leger P, Josse T, Blot G, Billecocq A, et al., A SAP30 complex inhibits IFN-beta expression in Rift Valley fever virus infected cells. PLoS Pathog, 2008. 4(1): p. e13. DOI: 10.1371/journal.ppat.0040013.

46. Leonard VH, Kohl A, Hart TJ, and Elliott RM, Interaction of Bunyamwera Orthobunyavirus NSs protein with mediator protein MED8: a mechanism for inhibiting the interferon response. J Virol, 2006. 80(19): p. 9667–75. DOI: 10.1128/JVI.00822-06.

47. Fitzmeyer EA, Dutt TS, Pinaud S, Graham B, Gallichotte EN, Hill JL, et al., A single-cell atlas of the Culex tarsalis midgut during West Nile virus infection. PLoS Pathog, 2025. 21(1): p. e1012855. DOI: 10.1371/journal.ppat.1012855.

48. Qu J, Betting V, van Iterson R, Kwaschik FM, and van Rij RP, Chromatin profiling identifies transcriptional readthrough as a conserved mechanism for piRNA biogenesis in mosquitoes. Cell Rep, 2023. 42(3): p. 112257. DOI: 10.1016/j.celrep.2023.112257.

49. Joo SS, Won TJ, Kim MJ, Hwang KW, and Lee DI, Interferon signal transduction of biphenyl dimethyl dicarboxylate/amantadine and anti-HBV activity in HepG2 2.2.15. Arch Pharm Res, 2006. 29(5): p. 405–11. DOI: 10.1007/BF02968591.

50. Hynes M, Ye W, Wang K, Stone D, Murone M, Sauvage F, et al., The seven-transmembrane receptor smoothened cell-autonomously induces multiple ventral cell types. Nat Neurosci, 2000. 3(1): p. 41–6. DOI: 10.1038/71114.

51. Ahearn YP, Saredy JJ, and Bowers DF, The Alphavirus Sindbis Infects Enteroendocrine Cells in the Midgut of Aedes aegypti. Viruses, 2020. 12(8). DOI: 10.3390/v12080848.

52. Hall DR, Johnson RM, Kwon H, Ferdous Z, Laredo-Tiscareno SV, Blitvich BJ, et al., Mosquito immune cells enhance dengue and Zika virus dissemination in Aedes aegypti. bioRxiv, 2024. DOI: 10.1101/2024.04.03.587950.

53. Ramirez L, Betanzos A, Raya-Sandino A, Gonzalez-Mariscal L, and Del Angel RM, Dengue virus enters and exits epithelial cells through both apical and basolateral surfaces and perturbs the apical junctional complex. Virus Res, 2018. 258: p. 39–49. DOI: 10.1016/j.virusres.2018.09.016.

54. Gerrard SR, Rollin PE, and Nichol ST, Bidirectional infection and release of Rift Valley fever virus in polarized epithelial cells. Virology, 2002. 301(2): p. 226–35. DOI: 10.1006/viro.2002.1588.

55. Singh VB, Singh MV, Gorantla S, Poluektova LY, and Maggirwar SB, Smoothened Agonist Reduces Human Immunodeficiency Virus Type-1-Induced Blood-Brain Barrier Breakdown in Humanized Mice. Sci Rep, 2016. 6: p. 26876. DOI: 10.1038/srep26876.

56. Doyle T, Moncorge O, Bonaventure B, Pollpeter D, Lussignol M, Tauziet M, et al., The interferon-inducible isoform of NCOA7 inhibits endosome-mediated viral entry. Nat Microbiol, 2018. 3(12): p. 1369–1376. DOI: 10.1038/s41564-018-0273-9.

57. Khan H, Winstone H, Jimenez-Guardeno JM, Graham C, Doores KJ, Goujon C, et al., TMPRSS2 promotes SARS-CoV-2 evasion from NCOA7-mediated restriction. PLoS Pathog, 2021. 17(11): p. e1009820. DOI: 10.1371/journal.ppat.1009820.

58. Yaple-Maresh ME, Flores GG, Zimmerman GE, Gomez-Rivera F, and Collins KL, Virus-induced vesicular acidification enhances HIV immune evasion. bioRxiv, 2024. DOI: 10.1101/2024.12.17.628989.

59. Petit MJ, Kenaston MW, Pham OH, Nagainis AA, Fishburn AT, and Shah PS, Nuclear dengue virus NS5 antagonizes expression of PAF1-dependent immune response genes. PLoS Pathog, 2021. 17(11): p. e1010100. DOI: 10.1371/journal.ppat.1010100.

60. Harmon B, Bird SW, Schudel BR, Hatch AV, Rasley A, and Negrete OA, A Genome-Wide RNA Interference Screen Identifies a Role for Wnt/beta-Catenin Signaling during Rift Valley Fever Virus Infection. J Virol, 2016. 90(16): p. 7084–7097. DOI: 10.1128/JVI.00543-16.

61. Ikegami T, Narayanan K, Won S, Kamitani W, Peters CJ, and Makino S, Dual functions of Rift Valley fever virus NSs protein: inhibition of host mRNA transcription and post-transcriptional downregulation of protein kinase PKR. Ann N Y Acad Sci, 2009. 1171 Suppl 1(Suppl 1): p. E75–85. DOI: 10.1111/j.1749-6632.2009.05054.x.

62. Billecocq A, Gauliard N, Le May N, Elliott RM, Flick R, and Bouloy M, RNA polymerase I-mediated expression of viral RNA for the rescue of infectious virulent and avirulent Rift Valley fever viruses. Virology, 2008. 378(2): p. 377–84. DOI: 10.1016/j.virol.2008.05.033.

63. Bonizzoni M, Dunn WA, Campbell CL, Olson KE, Dimon MT, Marinotti O, et al., RNA-seq analyses of blood-induced changes in gene expression in the mosquito vector species, Aedes aegypti. BMC Genomics, 2011. 12: p. 82. DOI: 10.1186/1471-2164-12-82.

64. Turell MJ, Wilson WC, and Bennett KE, Potential for North American mosquitoes (Diptera: Culicidae) to transmit rift valley fever virus. J Med Entomol, 2010. 47(5): p. 884–9. DOI: 10.1603/me10007.

65. Chowdhury A, Modahl CM, Tan ST, Wong Wei Xiang B, Misse D, Vial T, et al., JNK pathway restricts DENV2, ZIKV and CHIKV infection by activating complement and apoptosis in mosquito salivary glands. PLoS Pathog, 2020. 16(8): p. e1008754. DOI: 10.1371/journal.ppat.1008754.

66. Vera-Maloof FZ, Saavedra-Rodriguez K, Elizondo-Quiroga AE, Lozano-Fuentes S, and Black IV WC, Coevolution of the Ile1,016 and Cys1,534 Mutations in the Voltage Gated Sodium Channel Gene of *Aedes aegypti* in Mexico. PLoS Negl Trop Dis, 2015. 9(12): p. e0004263. DOI: 10.1371/journal.pntd.0004263.

67. Black WC and DuTeau NM, RAPD-PCR and SSCP analysis for insect population genetic studies, in The Molecular Biology of Insect Disease Vectors: A Methods Manual C.B.B.a.C.L. J. Crampton, Editor. 1997, Chapman and Hall: New York. p. 361–373.

68. Chen S, Ultrafast one-pass FASTQ data preprocessing, quality control, and deduplication using fastp. Imeta, 2023. 2(2): p. e107. DOI: 10.1002/imt2.107.

69. Carroll TS, Liang Z, Salama R, Stark R, and de Santiago I, Impact of artifact removal on ChIP quality metrics in ChIP-seq and ChIP-exo data. Front Genet, 2014. 5: p. 75. DOI: 10.3389/fgene.2014.00075.

70. Kim D, Paggi JM, Park C, Bennett C, and Salzberg SL, Graph-based genome alignment and genotyping with HISAT2 and HISAT-genotype. Nat Biotechnol, 2019. 37(8): p. 907–915. DOI: 10.1038/s41587-019-0201-4.

71. Matthews BJ, Dudchenko O, Kingan SB, Koren S, Antoshechkin I, Crawford JE, et al., Improved reference genome of Aedes aegypti informs arbovirus vector control. Nature, 2018. 563(7732): p. 501–507. DOI: 10.1038/s41586-018-0692-z.

72. Zhang Y, Liu T, Meyer CA, Eeckhoute J, Johnson DS, Bernstein BE, et al., Model-based analysis of ChIP-Seq (MACS). Genome Biol, 2008. 9(9): p. R137. DOI: 10.1186/gb-2008-9-9-r137.

73. Ramirez F, Ryan DP, Gruning B, Bhardwaj V, Kilpert F, Richter AS, et al., deepTools2: a next generation web server for deep-sequencing data analysis. Nucleic Acids Res, 2016. 44(W1): p. W160–5. DOI: 10.1093/nar/gkw257.

74. Ross-Innes CS, Stark R, Teschendorff AE, Holmes KA, Ali HR, Dunning MJ, et al., Differential oestrogen receptor binding is associated with clinical outcome in breast cancer. Nature, 2012. 481(7381): p. 389–93. DOI: 10.1038/nature10730.

75. Team RC, R: A language and environment for statistical computing. 2021, R Foundation for Statistical Computing: Vienna, Austria.

76. Sergushichev AA, An algorithm for fast preranked gene set enrichment analysis using cumulative statistic calculation. bioRxiv, 2016: p. 060012. DOI: 10.1101/060012.

77. Subramanian A, Tamayo P, Mootha VK, Mukherjee S, Ebert BL, Gillette MA, et al., Gene set enrichment analysis: a knowledge-based approach for interpreting genome-wide expression profiles. Proc Natl Acad Sci U S A, 2005. 102(43): p. 15545–50. DOI: 10.1073/pnas.0506580102.

78. Wang Q, Li M, Wu T, Zhan L, Li L, Chen M, et al., Exploring Epigenomic Datasets by ChIPseeker. Curr Protoc, 2022. 2(10): p. e585. DOI: 10.1002/cpz1.585.

79. Yu G, Wang LG, and He QY, ChIPseeker: an R/Bioconductor package for ChIP peak annotation, comparison and visualization. Bioinformatics, 2015. 31(14): p. 2382–3. DOI: 10.1093/bioinformatics/btv145.

80. Yu G and Gao C-H, enrichplot: Visualization of Functional Enrichment Result. 2025. p. R package version 1.28.4.

81. Wu T, Hu E, Xu S, Chen M, Guo P, Dai Z, et al., clusterProfiler 4.0: A universal enrichment tool for interpreting omics data. Innovation (Camb), 2021. 2(3): p. 100141. DOI: 10.1016/j.xinn.2021.100141.

82. Heinz S, Benner C, Spann N, Bertolino E, Lin YC, Laslo P, et al., Simple combinations of lineage-determining transcription factors prime cis-regulatory elements required for macrophage and B cell identities. Mol Cell, 2010. 38(4): p. 576–89. DOI: 10.1016/j.molcel.2010.05.004.

