## Supplementary material for "Altered histone modifications in *Aedes aegypti* following Rift Valley fever virus exposure": CutRun_ms_Suppl_Figs20250808.pptx

### Slide 1
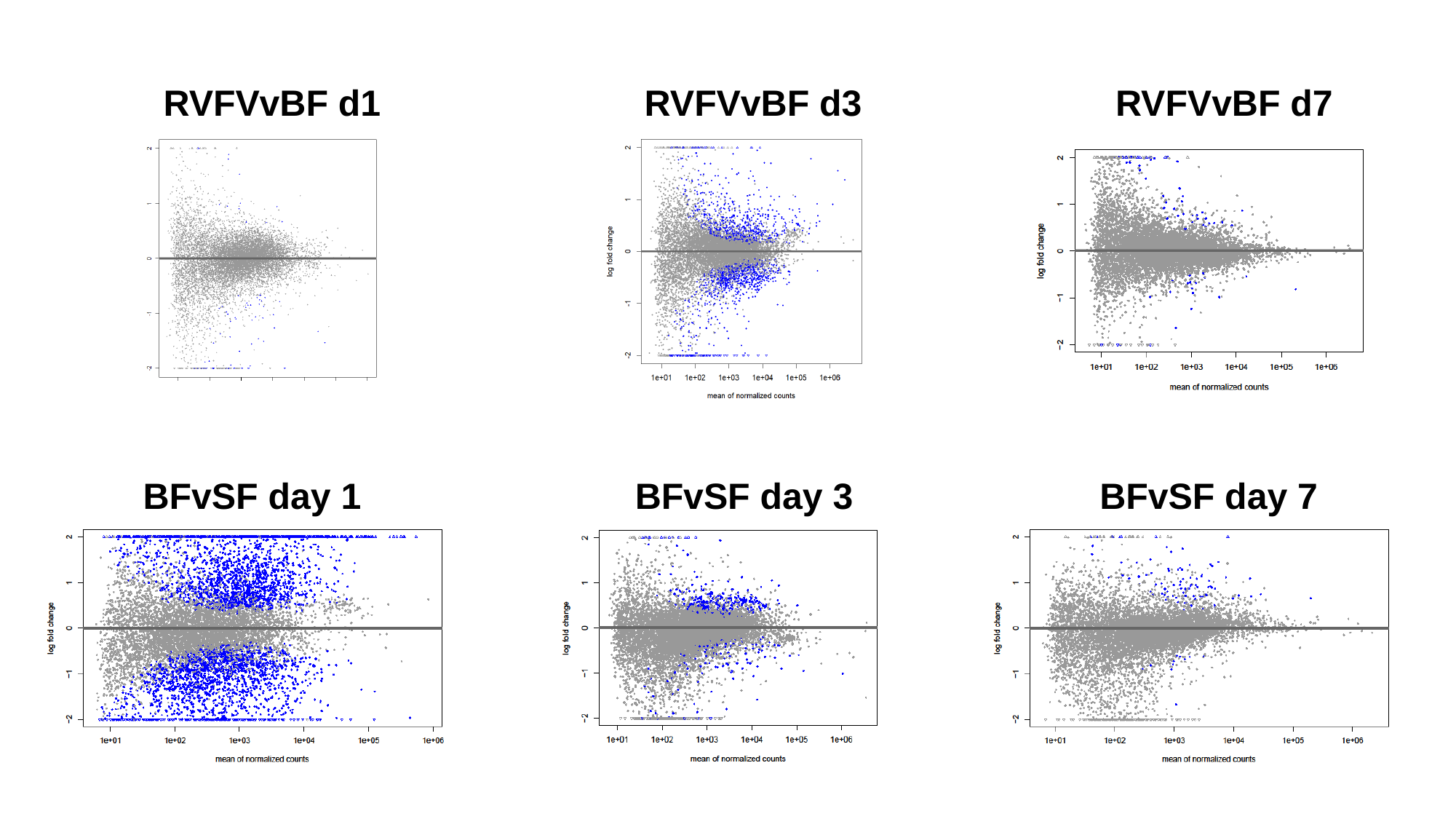

RVFVvBF d1 RVFVvBF d3 RVFVvBF d7
 BFvSF day 1 BFvSF day 3	 BFvSF day 7

### Slide 2
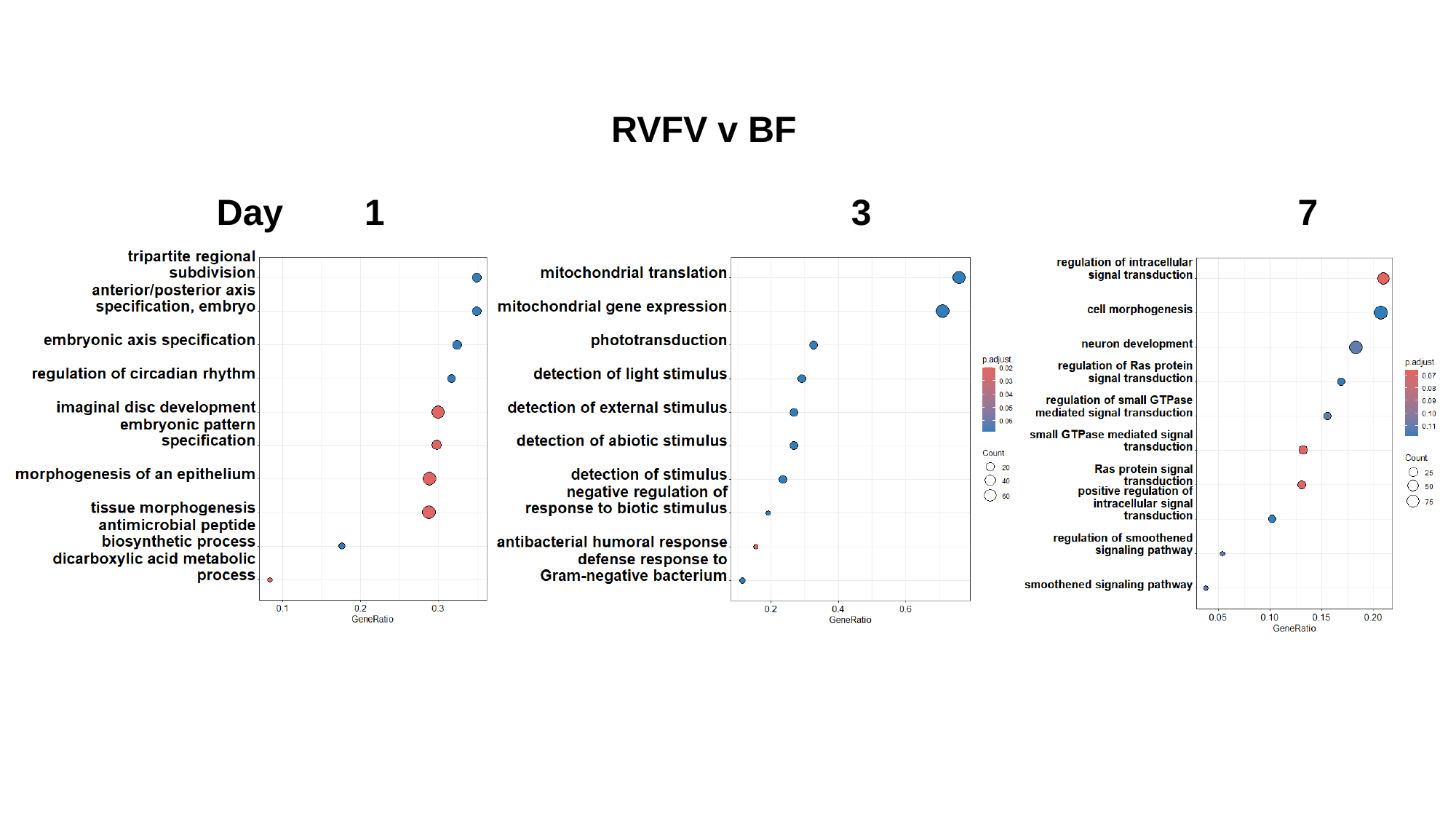

RVFV v BF
Day 1 3 7

### Slide 3
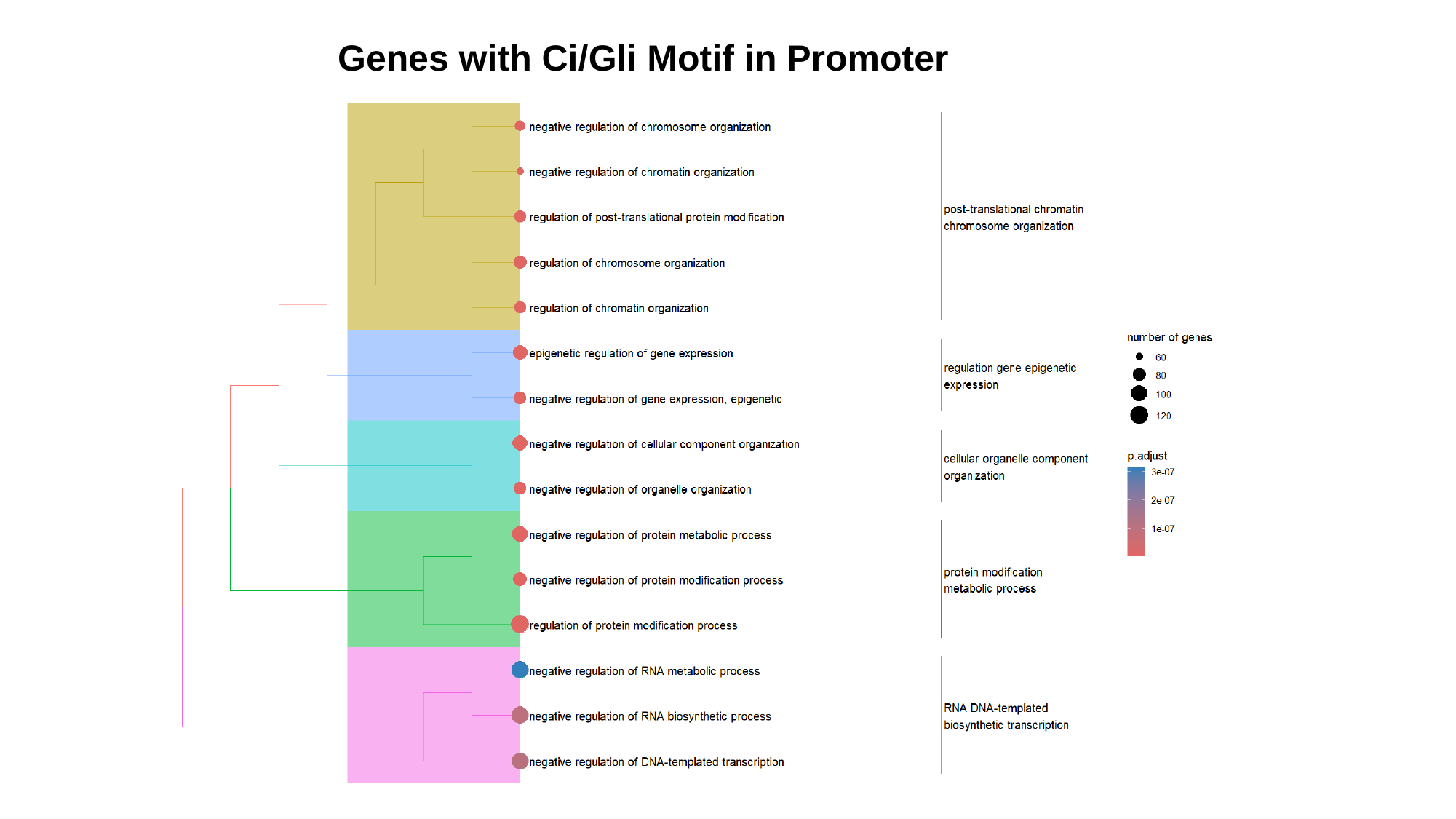

Genes with Ci/Gli Motif in Promoter

### Slide 4
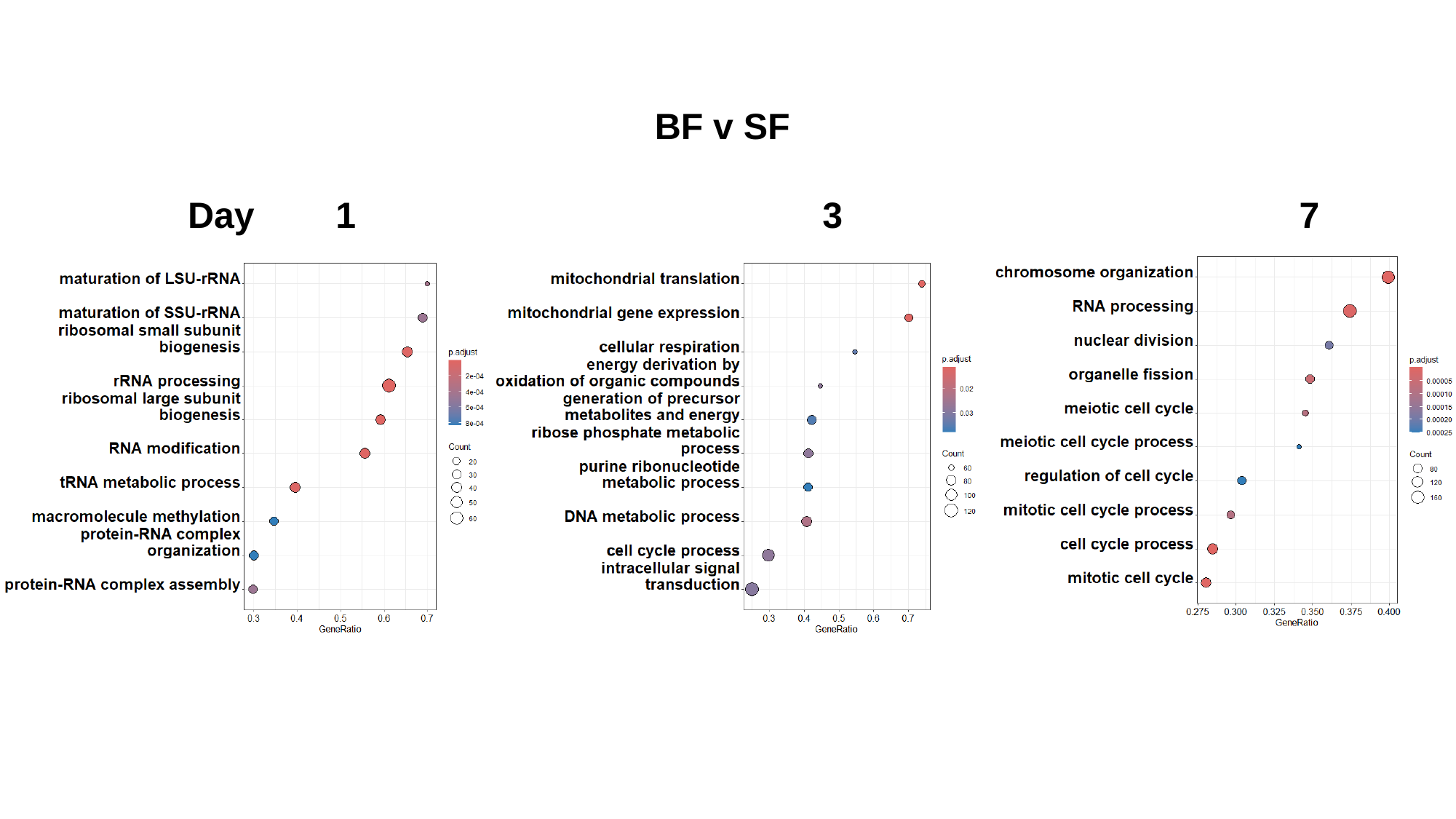

BF v SF
Day 1 3 7

### Slide 5
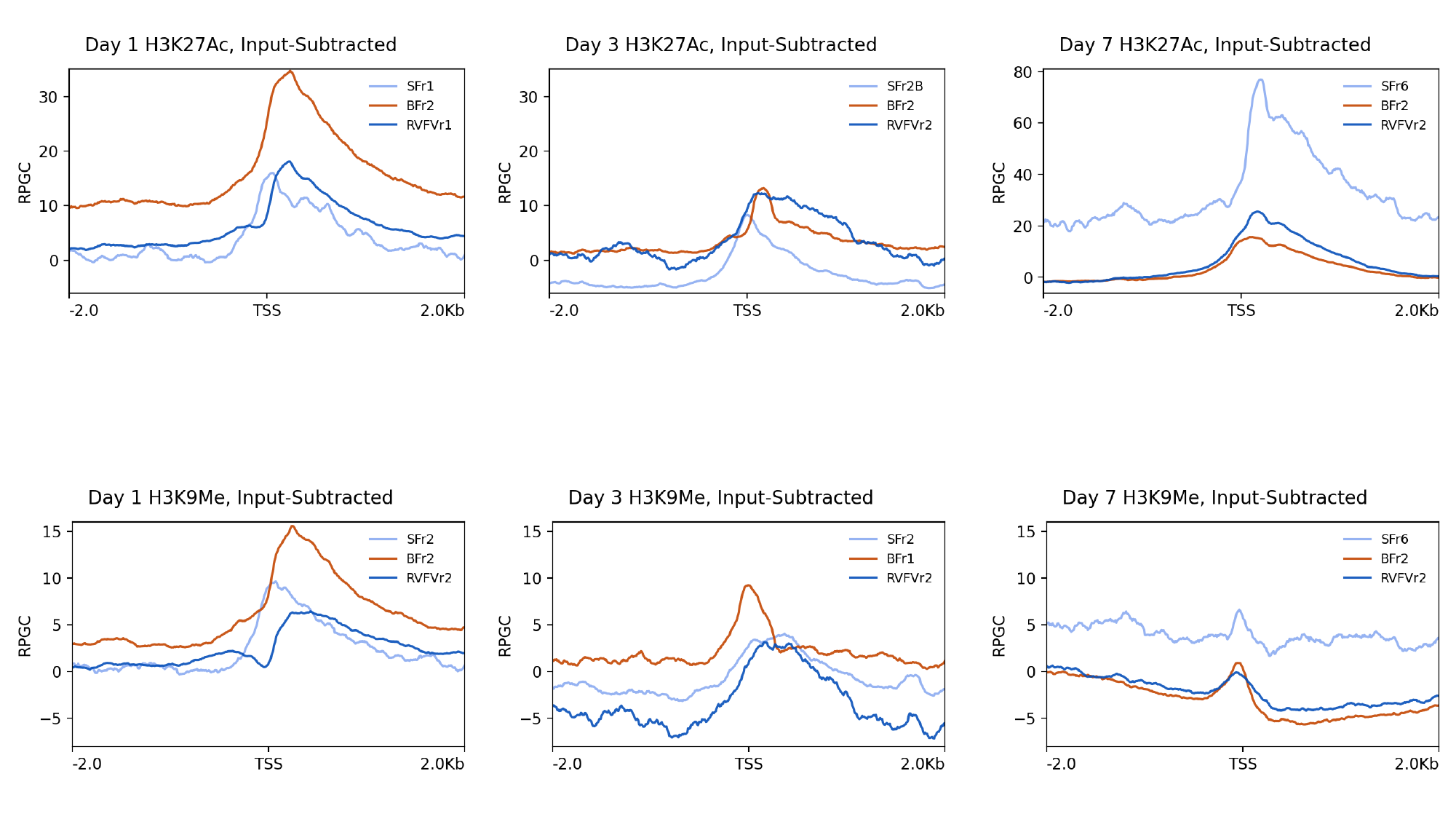

### Slide 6
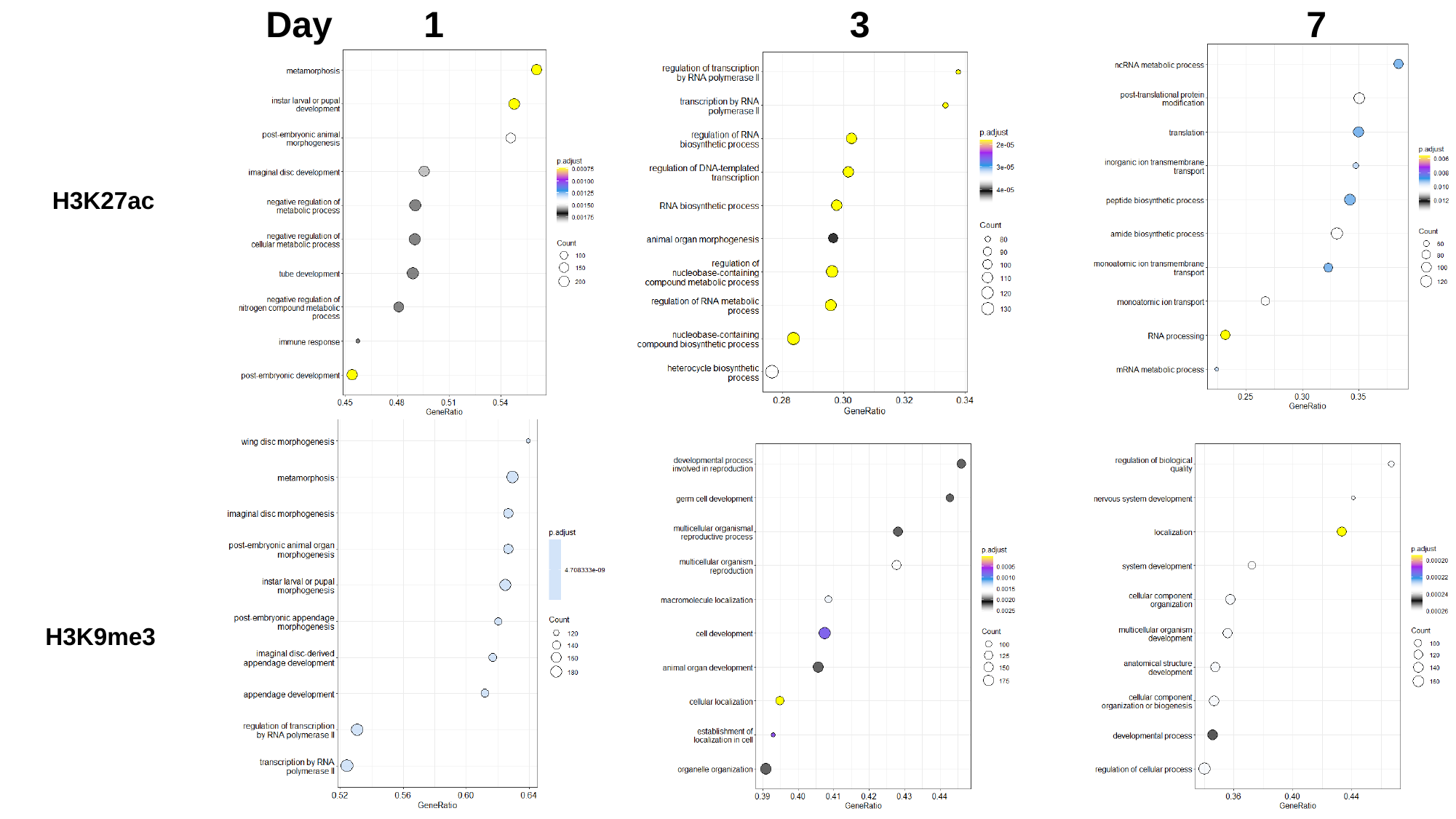

Day 1 3 7
 H3K27ac
 H3K9me3

### Slide 7
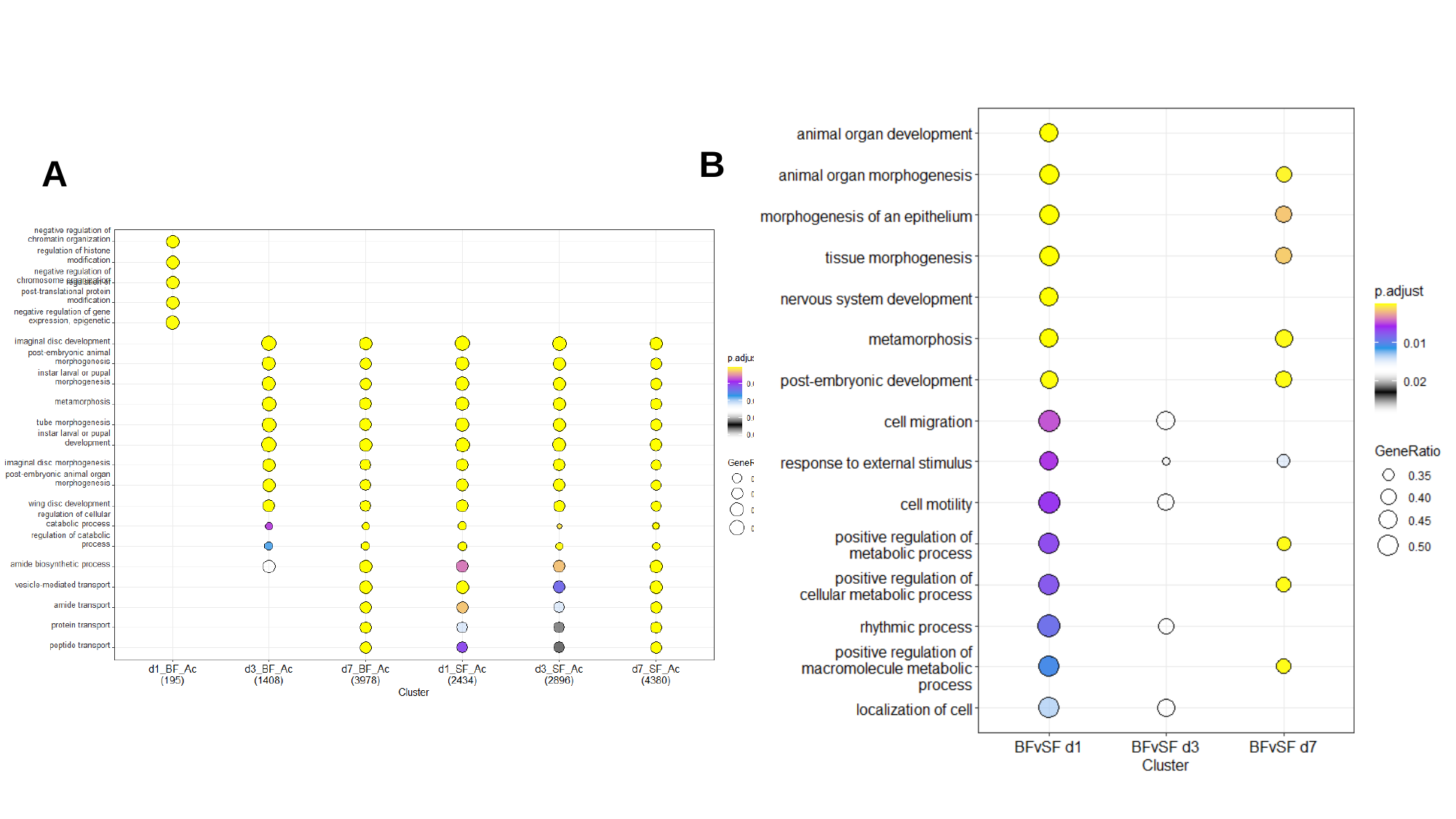

B
A

### Slide 8
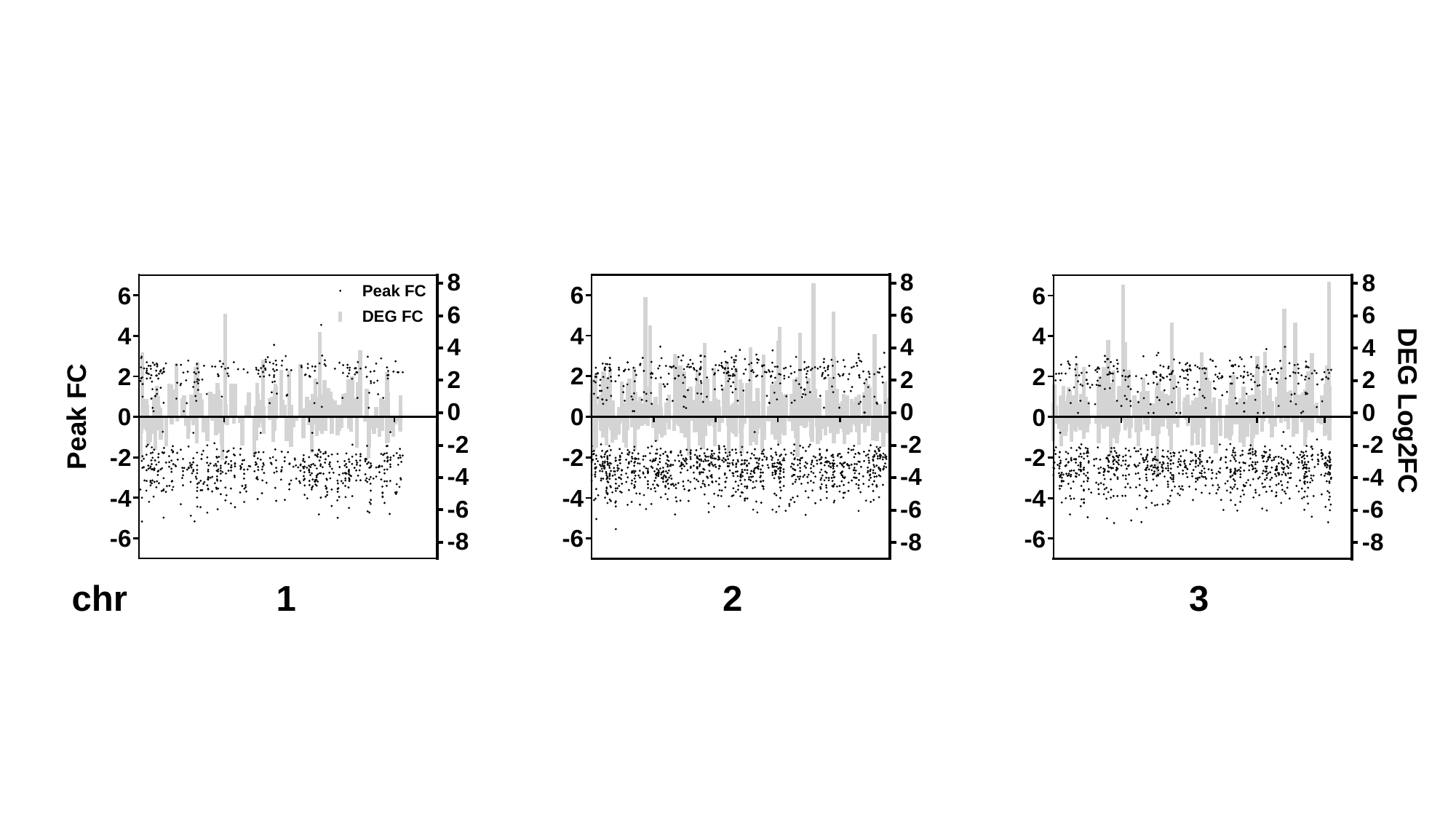

### Slide 9
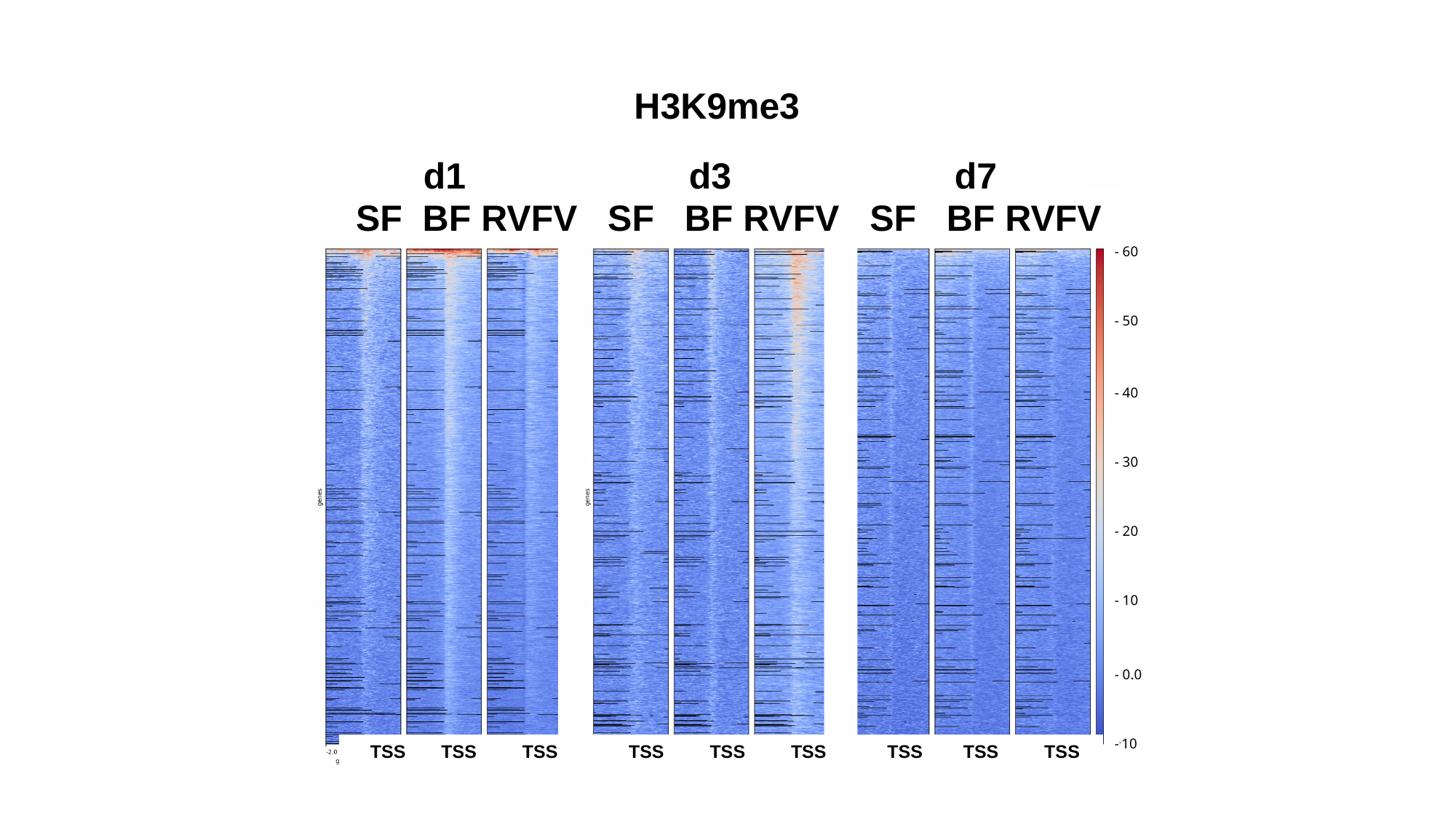

H3K9me3
H3K9me3
# d1 d3 d7
SF BF RVFV SF BF RVFV SF BF RVFV
- 60
- 50
- 40
- 30
- 20
- 10
- 0.0
-^-^10
 TSS TSS TSS TSS TSS TSS TSS TSS TSS

### Slide 10
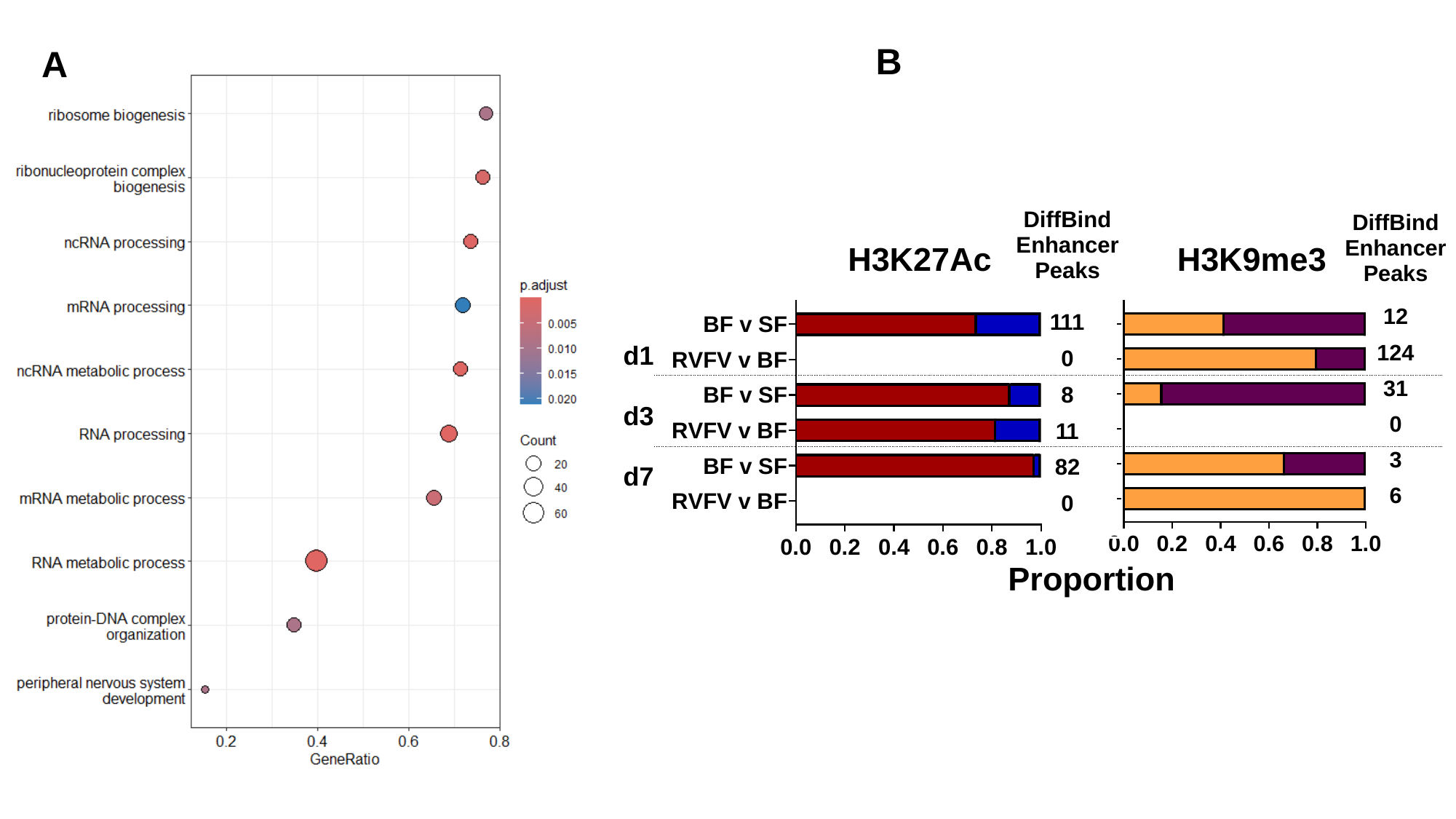

B
A
